# Similar enzymatic functions in distinct bioluminescence systems: Evolutionary recruitment of sulfotransferases in ostracod light organs

**DOI:** 10.1101/2023.04.12.536614

**Authors:** Emily S Lau, Jessica A Goodheart, Nolan T Anderson, Vannie L Liu, Arnab Mukherjee, Todd H Oakley

## Abstract

Genes from ancient families are sometimes involved in the convergent evolutionary origins of similar traits, even across vast phylogenetic distances. Sulfotransferases are an ancient family of enzymes that transfer sulfate from a donor to a wide variety of substrates, including probable roles in some bioluminescence systems. Here we demonstrate multiple sulfotransferases, highly expressed in light organs of the bioluminescent ostracod *Vargula tsujii*, transfer sulfate in vivo to the luciferin substrate, vargulin. We find luciferin sulfotransferases of ostracods are not orthologous to known luciferin sulfotransferases of fireflies or sea pansies; animals with distinct and convergently evolved bioluminescence systems compared to ostracods. Therefore, distantly related sulfotransferases were independently recruited at least three times, leading to parallel evolution of luciferin metabolism in three highly diverged organisms. Re-use of homologous genes is surprising in these bioluminescence systems because the other components, including luciferins and luciferases, are completely distinct. Whether convergently evolved traits incorporate ancient genes with similar functions or instead use distinct, often newer, genes may be constrained by how many genetic solutions exist for a particular function. When fewer solutions exist, as in genetic sulfation of small molecules, evolution may be more constrained to use the same genes time and again.

## Introduction

The convergent evolutionary origins of similar traits sometimes employ existing genetic elements that originated much earlier. This pattern of parallel evolution is often called “deep homology”, especially when similar but convergently evolved traits express shared transcription factors. For example, limbs in some distantly related phyla express the transcription factor *distalless* [1]. Convergently evolved traits also may recruit ancient genes with shared enzymatic functions. For example, convergently evolved instances of biomineralization use α-carbonic anhydrases, an ancient gene family found across animals [2] that facilitates conversion of carbon dioxide to bicarbonate, crucial in regulating pH and mineral deposition. Similarly, convergently evolved cases of photosymbiosis involve vacuolar H+-ATPase (*VHA*), another ancient gene family [3,4], which plays a vital role to acidify intracellular compartments, essential for nutrient transport and cellular metabolism. Finally, convergently evolved cases of bioluminescence may have recruited ancient dehalogenases [5,6]. Characterizing new examples where homology differs across levels of biological organization will build the knowledge necessary to answer fundamental evolutionary questions, such as what types of traits, genes, or evolutionary histories are more likely to lead to the use of ancient elements in novel systems [e.g. 7].

We hypothesize sulfotransferases, a family of enzymes shared across vast phylogenetic distances, are used in convergently evolved bioluminescence systems of distantly related species. Fireflies, sea pansies, and ostracods convergently evolved bioluminescence systems and produce light using structurally diverse organs or tissues [8–10], different luciferin substrates [11], and non-homologous, autogenic, luciferase enzymes [12,13] (figure S1). Despite these vast differences, these taxa use the same biochemical mechanism, sulfation — the transfer of sulfate (SO3-) from the donor 3′-phosphoadenosine-5′-phosphosulfate (PAPS) to a substrate [14] — to modulate the chemical state of luciferins [15–18] (figure S2). Sulfation may create chemical forms of the substrate that are better for storage by making them less susceptible to non-specific oxidation [17–20], or may create an activated form of the substrate for the luciferase reaction [21]. Sulfation is catalyzed by sulfotransferases in fireflies [17] and sea pansies [18], but less is known about ostracod sulfotransferases. Although previously published results do show crude protein extracts from the ostracod *Vargula hilgendorfii* can reversibly sulfate luciferin, thereby suggesting sulfation as a mechanism for luciferin metabolism in the organism [19], the genetic sequences of luciferin sulfotransferases in ostracods have yet to be identified and functionally characterized.

Here we provide evidence of independent recruitment of paralogous genes from a single, ancient gene family — sulfotransferases — leading to parallel evolution of luciferin sulfation in phylogenetically distant organisms with distinct bioluminescence systems. We use gene expression analyses, recombinant protein expression, and *in vitro* biochemical assays to identify and functionally test the five most highly expressed sulfotransferase genes from the light-producing organ of the cypridinid ostracod *Vargula tsujii* and report multiple sulfotransferases capable of sulfating vargulin *in vitro*. Our results, taken together with functional evidence of luciferin sulfotransferases previously published on sea pansies and fireflies [17,18] provide an example of an ancient and pervasive gene family that was recruited multiple times independently for a similar purpose.

## Materials and Methods

### *(a) Reference transcriptome assembly for* Vargula tsujii

We constructed a reference transcriptome for *Vargula tsujii* by pooling RNA from 88 specimens across various instars (see table S1; figure S3, S4A). We extracted RNA using TRIzol^(R)^ for sequencing at the BYU Sequencing Center using PacBio Sequel II (“IsoSeq”) using two transcript size fractions: 4-10 kb selected using BluePippin (Sage Science) and a non-size-selected fraction. Next, we processed circular consensus sequencing reads with the IsoSeq v3 pipeline (https://github.com/PacificBiosciences/IsoSeq) and combined them with Illumina short reads (SRR21201581, [22]) using rnaSPAdes with flags --rna, --only-assembler, and --trusted-contigs [23]. We clustered sequences with ≥ 95% identity using cd-hit-est [24,25].

### (b) Sulfotransferase candidates from the upper lip of Vargula tsujii

To quantify gene expression, we collected RNA from light organs (upper lips) of five male and four female ostracods maintained in culture [26] (figure S4B). After extracting RNA with Trizol, we used the Genomic Sequencing and Analysis Facility at UT-Austin for library preparations and Tag-Seq profiling using Illumina HiSeq 2500, SR100 [27] . We processed Tag-Seq reads (https://github.com/z0on/tag-based_RNAseq) by mapping to our reference transcriptome with Bowtie2 v.2.3.4.3 [28] (table S2). For protein sequence identification, we translated sequences with TransDecoder v.5.5.0 and used HMMER v.3.2.1 (hmmer.org) to find complete protein sequences containing a sulfotransferase domain (Pfam ID: PF00685) of e-value ≤ 1e-3.

### (c) Cloning, expression, and purification of Vargula tsujii sulfotransferase candidates

We synthesized DNA sequences for candidate luciferin sulfotransferases ST1-5 (figure S5) as gBlocks™ (Integrated DNA Technologies) and cloned them into bacterial expression vectors (ST3 and ST4 into pQE80L; ST1, ST2, and ST5 in pET SUMO to improve solubility [29] (figure S6, figure S7) using Gibson assembly (table S3). These vectors allow protein expression with N-terminal hexahistidine tags for purification using metal-affinity chromatography. We propagated plasmid constructs in *E. coli* NEB^®^ 10-beta cells and verified by Sanger sequencing (Genewiz, South Plainfield, NJ).

We expressed proteins in *E. coli* BL21 cells via induction with IPTG, followed by incubating cultures at room temperature for 18 hours. We purified proteins by incubating lysed cell extracts with Ni-NTA agarose beads, then using gravity-flow metal-affinity chromatography to elute hexahistidine-tagged proteins. We concentrated eluates by spin filtration (10 kDa molecular weight cutoff) and assessed protein purity via SDS-PAGE (figure S7, table S4) and densitometry. We quantified purified protein yields using a Bradford assay (Bio-Rad Laboratories, Hercules, CA) calibrated with a bovine serum albumin standard. These methods successfully purified all five proteins with > 90% homogeneity (table S5), and we prepared proteins freshly for each experiment.

### (d) Sulfotransferase Functional Assays

We tested for sulfotransferase activity using two functional assays. First, we used a commercially available sulfotransferase assay (R&D Systems, Minneapolis, MN) that probes for the formation of 3′-phosphoadenosine-5′-phosphate (PAP), a byproduct of sulfation, by measuring the increase in absorbance at 620 nm following the enzymatic hydrolysis of PAP into phosphate, which binds to malachite green. In this assay, the phosphatase used to hydrolyze PAP into phosphate has some activity with PAPS, so we used an enzyme-free negative control and kept PAPS the same between experiments and control.

The second assay quantifies decreases in luminescence to test each candidate protein’s ability to convert luciferin to its sulfated form, which luciferase does not oxidize. For ostracod luciferase, we transfected CHO cells with a pCMV *Cypridina* luciferase vector, and purified the protein with metal-affinity chromatography (figure S8). We first mixed each purified protein with ostracod luciferin and PAPS, incubated at 25 °C for 3 hours, then quenched reactions by heat-inactivation. After cooling, we added the diluted reaction product to purified ostracod luciferase and immediately measured bioluminescence. We compared luminescence between test and enzyme-free reactions with a decrease in light emission indicating less luciferin available for light production due to conversion to its sulfated form. In addition to luciferin sulfation assays, we used an established p-nitrophenol sulfation assay to evaluate sulfotransferase candidates for their ability to sulfate p-nitrophenol, a common substrate for many sulfotransferases, including those unrelated to bioluminescence (figure S9).

### (e) Phylogenetic analyses

We estimated a phylogenetic tree for sulfotransferase candidates from *V. tsujii* and a tree of functionally tested sulfotransferases from the literature (as of March 2023) (table S6, Supplemental References. For aligning protein sequences, we used MAFFT (v. 7.429) [30], trimmed the alignment of functionally tested sulfotransferases using trimAl (v.1.4) [31] with the flag -gappyout, inferred maximum likelihood phylogenies using IQ-TREE (v. 1.6.12) [32], assuming the best-fit substitution model (LG+I+G4 for candidates, LG+F+R5 for functionally tested, LG+R5 for trimmed alignment of functionally tested) according to Bayesian Information Criterion, as determined with ModelFinder [33], and performed ultrafast bootstrap approximation [34] with 1000 bootstrap replicates. To midpoint root the phylogeny and visualize bootstrap values and expression values, we used the R packages phytools (v. 1.5-1) [35] and ggtree (v. 3.9.0) [36].

## Results

### *Functional assays of sulfotransferase genes from the light organ of* Vargula tsujii

From Tag-seq on light-producing organs of *Vargula tsujii* (N = 9), we identified 40 genes, expressed in at least one individual, that contain domains with significant similarity to sulfotransferase domains, with nine expressed in all individuals. We selected the five (ST1-5) most highly expressed candidate sulfotransferases in light organs (figure S10) for recombinant expression and performed two functional assays to test for luciferin (vargulin) sulfation activity (figure 1b,c). Based on the commercial absorbance-based assay that probes for sulfotransferase-dependent conversion of PAPS to PAP, we found three candidates have the highest fold-change (5-6 fold) in activity (ST3, ST4, ST5) with ST1 also showing statistically significant change in absorbance under the conditions used (figure 1b). Based on the bioluminescence assay, which measures the decrease in light emission caused by sulfating luciferin, we infer the largest magnitude of luciferin sulfation in ST3 and ST5 (figure 1c).

**Figure 1.**
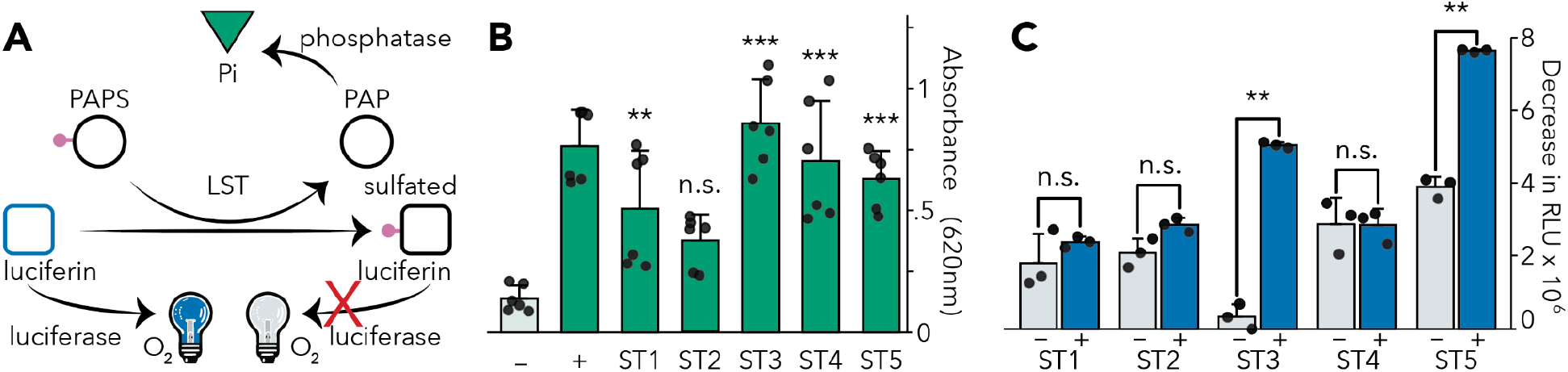
Based on two different assays, multiple candidate sulfotransferases from *Vargula tsujii*, especially ST3 and ST5, exhibit luciferin sulfation activity. **A**) Although mass spectroscopy would be the most direct demonstration of vargulin sulfation, we illustrate a schematic of the chemistry underlying the two assays we use here. **B**) Results of the absorbance-based sulfotransferase assay that probes for the conversion of PAPS (sulfate donor) to PAP. Our positive (+) control used 50 μL of 10 ng/μL Golgi-resident PAP-specific 3’ phosphatase and 50 μM PAP to confirm phosphatase activity and a negative (–) control used 50 μL of 10 ng/μL coupling phosphatase, 100 μM luciferin, 200 μM lithium-free PAPS, but lacks any sulfotransferase. The test reactions used 50 μL of 10 ng/μL coupling phosphatase, 100 μM luciferin, and 200 μM lithium-free PAPS, and 10 μM sulfotransferase. Three genes showed the highest activity (ST3-5) and ST1 also showed significant activity under these conditions. **C**) Results of the bioluminescence assay that probes for a decrease in light emission due to sulfation of luciferin, which decreases luciferin available for oxidation. Negative control reactions (gray bars) lack PAPS. Test reactions (blue bars) included 10 μL of 10 μM ostracod luciferin and 100 uM PAPS with 1 μL of each purified protein (final concentrations ST1: 15 μM, ST2: 22 μM, ST3: 51 μM, ST4: 32 μM, ST5: 25 μM). Variations across the negative controls likely arise from the spontaneous oxidation of luciferin. Compared to control PAPS-free reactions, two candidates (ST3, ST5) exhibit a significant decrease in light production indicative of sulfation activity sufficient to modulate bioluminescence in vitro. ST2 may have some activity, but was not statistically significant with Welch’s test.

### Ostracods, fireflies, and sea pansies sulfate luciferin by using homologous genes

Genes of ostracods, fireflies, and sea pansies convergently evolved sulfation capabilities (figure 2), probably independently recruiting sulfotransferase genes as a mechanism for metabolizing luciferin. Our phylogenetic analysis of functionally characterized luciferin sulfotransferases (LSTs) are most closely related to sulfotransferases of non-luminous animals. We find firefly LST to be most closely related to a gene from a non-luminous silk moth *Bombyx mori* that sulfates multiple substrates. The LST of the sea pansy *R. muelleri* is most closely related to sulfotransferase genes from a non-luminous tick (*Ixodes scapularis*) and a nematode (*Caenorhabditis elegans*). The five candidate LSTs from the ostracod *V. tsujii* we tested form a clade sister to genes from non-luminous *B. mori*, and *D. melanogaster*, and ST3 from firefly, which does not sulfate luciferin [17]. The close relationship between luciferin sulfotransferases to genes from non-luminous animals or genes lacking the ability to sulfate luciferin, strongly support convergent recruitment of LSTs in fireflies, sea pansies, and ostracods (figure 2, figure S11).

**Figure 2.**
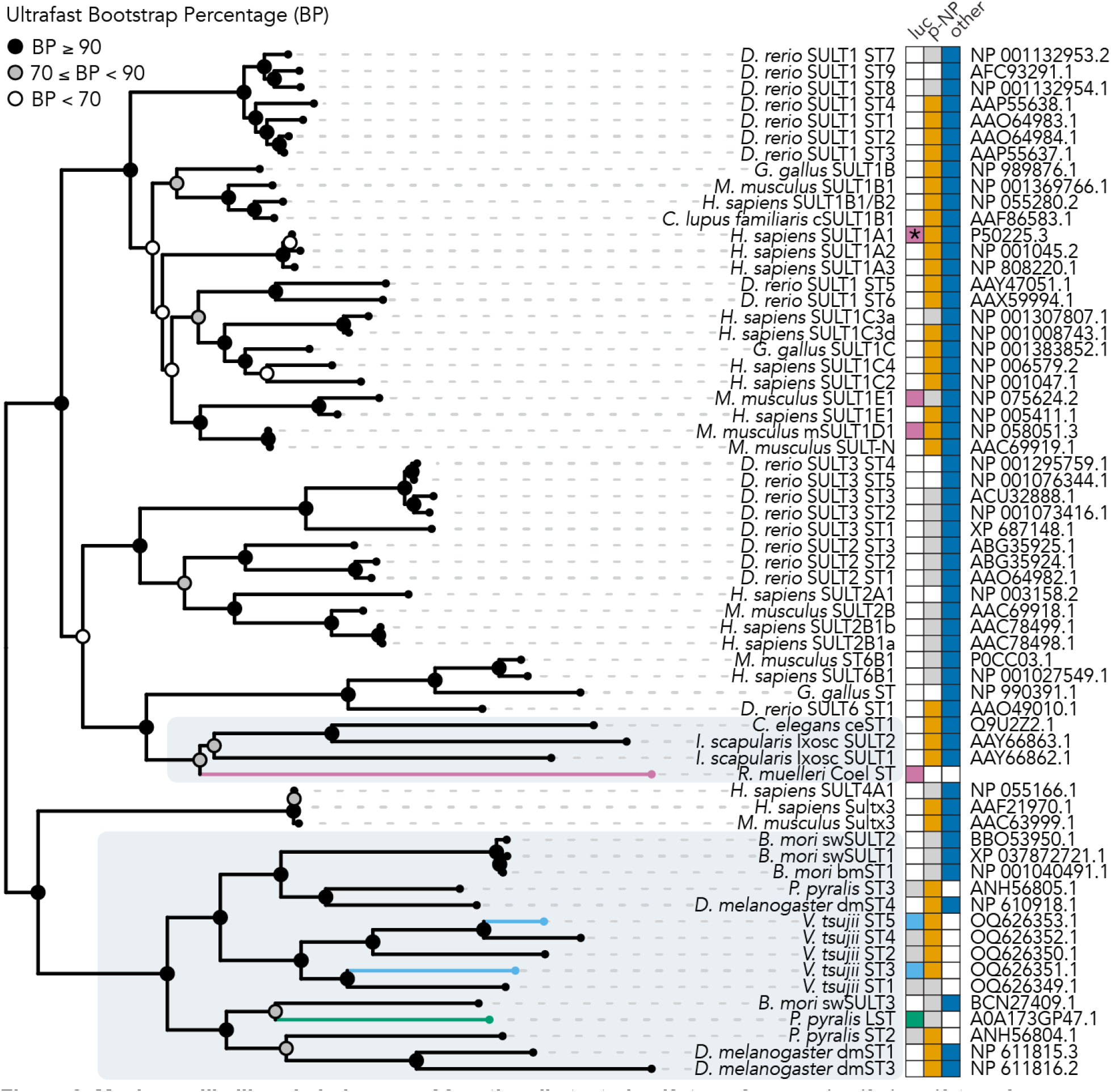
Maximum-likelihood phylogeny of functionally tested sulfotransferases. Luciferin sulfotransferases from luminous organisms are denoted by colored branches (firefly = green, ostracods = light blue, sea pansies = purple). Invertebrate sulfotransferases are highlighted in gray. Colored squares (green = firefly luciferin, light blue = ostracod luciferin for genes with strong activity in both assays herein, purple = sea pansy luciferin, orange = p-nitrophenol, dark blue = other) indicate functionally demonstrated substrates for each sulfotransferase; gray squares indicate molecules that are not sulfated by the respective sulfotransferase; white squares are untested. *Sulfates the luciferin coelenterazine at a different site compared to sea pansy sulfated luciferin [37]. Accession numbers are listed on the right. Ultrafast bootstrap values are represented by colored circles at each node.

## Discussion

Convergently evolved traits may independently deploy members of deeply conserved gene families, a pattern called deep homology [1]. Although criticized for imprecision and incompleteness [38,39], the term deep homology highlights the importance of ancient evolutionary histories in shaping new traits [40]. Here we report sulfation in bioluminescence systems may have evolved by independently recruiting homologous genes from the ancient sulfotransferase gene family present across animals. We provide functional evidence for multiple sulfotransferases capable of sulfating luciferin in the ostracod *Vargula tsujii* and synthesize our results with those previously described in fireflies and sea pansies.

Although establishing the genetic basis of organismal functions requires gene knockdowns, multiple lines of evidence point to sulfotransferase involvement in the bioluminescence of ostracods. Our results demonstrate multiple sulfotransferase genes of *V. tsujii* catalyze a reaction to sulfate luciferin, but determining how many using statistical significance depends on the assay (figure 1). The commercial assay that probes for PAP indicates four genes with statistically significant sulfation of luciferin (with ST3-5 showing especially strong differences between test and control) and the ‘bioluminescence assay’ that probes directly for luciferin sulfation indicates two genes (ST3 and ST5). The statistical difference between assays is likely due to specific experimental conditions, with the bioluminescence assay requiring a lower concentration of luciferin (10 μM) than the absorbance assay (100 μM), due to substrate inhibition of luciferase (figure S12). Nevertheless, results from both assays support multiple sulfotransferases highly expressed in the light organ — especially ST3 and ST5 — are capable of sulfating luciferin. Forthcoming work indicates ST3 is the only sulfotransferase with significant co-expression with luciferase across multiple stages and tissues (Oakley, In Prep). Based on this co-expression with luciferase, we speculate ST3 could be the primary sulfotransferase used in bioluminescence. Coupled with published results from crude protein extracts of *V. hilgendorfii* that sulfate luciferin [19], we hypothesize ST3 and possibly at least one other sulfotransferase (ST5) have important organismal functions in luminous ostracods, perhaps for use as a storage form of the easily oxidized luciferin.

A similar confluence of evidence suggests sulfotransferases are important to bioluminescence systems of fireflies and sea pansies. In fireflies, one sulfotransferase, highly expressed in lanterns, can catalyze formation of sulfoluciferin from luciferin and the reverse reaction [17]. This suggests an organismal function for a luciferin sulfotransferase in fireflies, again perhaps for creating a storage form of the substrate [17]. In the anthozoan *R. muelleri*, sulfated luciferin is converted into luciferin, bound to a luciferin binding protein, and released upon addition of calcium to react with luciferase [41]. Recently, two sulfotransferase isoforms, similar in sequence to the native protein, were cloned, expressed, and shown to have sulfotransferase activity [18]. Because preparations from *Renilla* yield much higher amounts of luciferin sulfate compared to the amount of luciferin bound to luciferin binding proteins [42], the sulfated form probably acts as a storage form of the luminous substrate in the animal [43]. Therefore, despite difficulties in knocking down specific sulfotransferases in ostracods, fireflies, or sea pansies, the combination of native preparations, gene expression studies, and heterologous expression suggest these animals use sulfotransferases (figure S13) in their respective bioluminescence systems.

Despite a growing realization of deep homology in evolutionary history, discovering homologous components in bioluminescence systems is arresting because of the distinctness of the other components. First, even the small-molecule substrates catalyzed by sulfotransferases are different: the luciferin of ostracods is vargulin, fireflies is D-luciferin, and sea pansy is coelenterazine. The luciferases also evolved from non-homologous gene families [12,13,44]. At the organismal level, bioluminescence is created by very different structures. Ostracod bioluminescence is secreted outside the body by the upper lip [45], firefly light is created in an abdominal structure called the lantern, and in sea pansies, light production occurs in specific cells called photocytes, located in the endoderm of two types of polyps. The presence of homologous sulfotransferases with similar functions along with non-homologous luciferases and distinct substrates provides a striking example of the often varied, cobbled-together elements of evolved complex systems. As more examples accumulate of cases of ancient genes independently recruited for similar functions, a logical pattern begins to emerge. Deeply homologous genes tend to have generally useful functions, such as sulfating small molecules in the case of sulfotransferases. Especially when there are few other solutions — as there is no other demonstrated mechanism for sulfation besides sulfotransferases – evolution may recruit the same gene family time and again, even in animals as distantly related as cnidarians and arthropods. This highlights how constraints — in this case determined by the number of biological solutions — may bias the options available to evolution [46,47].

## Author Contributions

ESL: Conceptualization, Methodology, Software, Formal analysis, Investigation, Data Curation, Writing - Original Draft and Review & Editing, Visualization. THO: Conceptualization, Resources, Writing - Original Draft and Review & Editing, Supervision, Project administration, Funding acquisition. AM: Methodology, Resources, Writing - Review & Editing, Supervision, Project administration, Funding acquisition. JAG: Software, Formal analysis, Investigation, Writing - Review & Editing, Resources, Data Curation. NTA: Methodology, Investigation. VLL: Investigation.

## Data Availability

All raw sequencing data are available on NCBI (BioProject: PRJNA935772). Sulfotransferase sequences are available on GenBank (Accessions: OQ626349, OQ626350, OQ626351, OQ626352, OQ626353). Plasmid sequences, code for analyzing and visualizing data, and data files for gene expression, phylogenetic analyses, and functional assays are available on Dryad (https://doi.org/10.25349/D94W5N).

## References Cited

1. Shubin N, Tabin C, Carroll S. 2009 Deep homology and the origins of evolutionary novelty. Nature 457, 818–823.

2. Le Roy N, Jackson DJ, Marie B, Ramos-Silva P, Marin F. 2014 The evolution of metazoan α-carbonic anhydrases and their roles in calcium carbonate biomineralization. Front. Zool. 11, 1–16.

3. Barott KL, Venn AA, Perez SO, Tambutté S, Tresguerres M. 2015 Coral host cells acidify symbiotic algal microenvironment to promote photosynthesis. Proc. Natl. Acad. Sci. U. S. A. 112, 607–612.

4. Armstrong EJ, Roa JN, Stillman JH, Tresguerres M. 2018 Symbiont photosynthesis in giant clams is promoted by V-type H-ATPase from host cells. J. Exp. Biol. 221. (doi:10.1242/jeb.177220)

5. Delroisse J, Ullrich-Lüter E, Blaue S, Ortega-Martinez O, Eeckhaut I, Flammang P, Mallefet J. 2017 A puzzling homology: a brittle star using a putative cnidarian-type luciferase for bioluminescence. Open Biol. 7. (doi:10.1098/rsob.160300)

6. Chaloupkova R et al. 2019 Light-Emitting Dehalogenases: Reconstruction of Multifunctional Biocatalysts. ACS Catal. 9, 4810–4823.

7. Lau ES, Oakley TH. 2021 Multi-level convergence of complex traits and the evolution of bioluminescence. Biol. Rev. Camb. Philos. Soc. 96, 673–691.

8. Peterson MK, Buck J. 1968 Light organ fine structure in certain Asiatic fireflies. Biol. Bull. 135, 335–348.

9. Spurlock BO, Cormier MJ. 1975 A fine structure study of the anthocodium in Renilla mülleri: Evidence for the existence of a bioluminescent organelle, the luminelle. J. Cell Biol. 64, 15–28.

10. Huvard AL. 1993 Ultrastructure of the light organ and immunocytochemical localization of luciferase in luminescent marine ostracods (Crustacea: Ostracoda: Cypridinidae). J. Morphol. 218, 181–193.

11. Tsarkova AS. 2021 Luciferins Under Construction: A Review of Known Biosynthetic Pathways. Frontiers in Ecology and Evolution 9. (doi:10.3389/fevo.2021.667829)

12. Lorenz WW, McCann RO, Longiaru M, Cormier MJ. 1991 Isolation and expression of a cDNA encoding Renilla reniformis luciferase. Proc. Natl. Acad. Sci. U. S. A. 88, 4438–4442.

13. Oba Y, Ojika M, Inouye S. 2003 Firefly luciferase is a bifunctional enzyme: ATP-dependent monooxygenase and a long chain fatty acyl-CoA synthetase. FEBS Lett. 540, 251–254.

14. Falany CN. 1997 Enzymology of human cytosolic sulfotransferases. FASEB J. 11, 206–216.

15. Hori K, Nakano Y, Cormier MJ. 1972 Studies on the bioluminescence of Renilla reniformis. XI. Location of the sulfate group in luciferyl sulfate. Biochim. Biophys. Acta 256, 638–644.

16. Nakamura M, Suzuki T, Ishizaka N, Sato J-I. 2014 Identification of 3-enol sulfate of Cypridina luciferin, Cypridina luciferyl sulfate, in the sea-firefly Cypridina (Vargula) hilgendorfii. Tetrahedron 70, 2161–2168.

17. Fallon TR, Li F-S, Vicent MA, Weng J-K. 2016 Sulfoluciferin is Biosynthesized by a Specialized Luciferin Sulfotransferase in Fireflies. Biochemistry 55, 3341–3344.

18. Tzertzinis G, Baker B, Benner J, Brown E, Corrêa IR Jr, Ettwiller L, McClung C, Schildkraut I. 2022 Coelenterazine sulfotransferase from Renilla muelleri. PLoS One 17, e0276315.

19. Nakamura M, Suzuki T, Ishizaka N, Sato J-I, Inouye S. 2014 Identification of 3-enol sulfate of Cypridina luciferin, Cypridina luciferyl sulfate, in the sea-firefly Cypridina (Vargula) hilgendorfii. Tetrahedron 70, 2161–2168.

20. Guo Y-J, Cui C-X, Liu Y-J. 2022 Theoretical Study on Storage and Release of Firefly Luciferin. Photochem. Photobiol. 98, 184–192.

21. Inoue S, Kakoi H, Goto T. 1976 Squid bioluminescence III. Isolation and structure of Watasenia luciferin. Tetrahedron Lett. 17, 2971–2974.

22. Ellis EA et al. 2022 Sexual signals persist over deep time: ancient co-option of bioluminescence for courtship displays in cypridinid ostracods. Syst. Biol. (doi:10.1093/sysbio/syac057)

23. Bushmanova E, Antipov D, Lapidus A, Prjibelski AD. 2019 rnaSPAdes: a de novo transcriptome assembler and its application to RNA-Seq data. Gigascience 8. (doi:10.1093/gigascience/giz100)

24. Li W, Godzik A. 2006 Cd-hit: a fast program for clustering and comparing large sets of protein or nucleotide sequences. Bioinformatics 22, 1658–1659.

25. Fu L, Niu B, Zhu Z, Wu S, Li W. 2012 CD-HIT: accelerated for clustering the next-generation sequencing data. Bioinformatics 28, 3150–3152.

26. Goodheart JA et al. 2020 Laboratory culture of the California Sea Firefly Vargula tsujii (Ostracoda: Cypridinidae): Developing a model system for the evolution of marine bioluminescence. Sci. Rep. 10, 10443.

27. Meyer E, Aglyamova GV, Matz MV. 2011 Profiling gene expression responses of coral larvae (Acropora millepora) to elevated temperature and settlement inducers using a novel RNA-Seq procedure. Mol. Ecol. 20, 3599–3616.

28. Langmead B, Salzberg SL. 2012 Fast gapped-read alignment with Bowtie 2. Nat. Methods 9, 357–359.

29. Butt TR, Edavettal SC, Hall JP, Mattern MR. 2005 SUMO fusion technology for difficult-to-express proteins. Protein Expr. Purif. 43, 1–9.

30. Katoh K, Standley DM. 2013 MAFFT multiple sequence alignment software version 7: improvements in performance and usability. Mol. Biol. Evol. 30, 772–780.

31. Capella-Gutiérrez S, Silla-Martínez JM, Gabaldón T. 2009 trimAl: a tool for automated alignment trimming in large-scale phylogenetic analyses. Bioinformatics 25, 1972–1973.

32. Nguyen L-T, Schmidt HA, von Haeseler A, Minh BQ. 2015 IQ-TREE: a fast and effective stochastic algorithm for estimating maximum-likelihood phylogenies. Mol. Biol. Evol. 32, 268–274.

33. Kalyaanamoorthy S, Minh BQ, Wong TKF, von Haeseler A, Jermiin LS. 2017 ModelFinder: fast model selection for accurate phylogenetic estimates. Nat. Methods 14, 587–589.

34. Hoang DT, Chernomor O, von Haeseler A, Minh BQ, Vinh LS. 2017 UFBoot2: Improving the Ultrafast Bootstrap Approximation. Mol. Biol. Evol. 35, 518–522.

35. Revell LJ. 2012 phytools: an R package for phylogenetic comparative biology (and other things). Methods Ecol. Evol. 3, 217–223.

36. Yu G, Smith DK, Zhu H, Guan Y, Lam TT-Y. 2017 Ggtree: An r package for visualization and annotation of phylogenetic trees with their covariates and other associated data. Methods Ecol. Evol. 8, 28–36.

37. Inouye S, Matsuda K, Nakamura M. 2023 Enzymatic sulfation of coelenterazine by human cytosolic aryl sulfotransferase SULT1A1: identification of coelenterazine C2-benzyl monosulfate by LC/ESI-TOF-MS. Biochem. Biophys. Res. Commun. 665, 133–140.

38. Scotland RW. 2010 Deep homology: a view from systematics. Bioessays 32, 438–449.

39. DiFrisco J, Wagner GP, Love AC. 2023 Reframing research on evolutionary novelty and co-option: Character identity mechanisms versus deep homology. Semin. Cell Dev. Biol. 145, 3–12.

40. Rose MR, Oakley TH. 2007 The new biology: beyond the Modern Synthesis. Biol. Direct 2, 30.

41. Anderson JM, Charbonneau H, Cormier MJ. 1974 Mechanism of calcium induction of Renilla bioluminescence. Involvement of a calcium-triggered luciferin binding protein. Biochemistry 13, 1195–1200.

42. Shimomura O, Johnson FH. 1979 Comparison of the amounts of key components in the bioluminescence systems of various coelenterates. Comparative Biochemistry and Physiology Part B: Comparative Biochemistry 64, 105–107.

43. Cormier MJ, Hori K, Karkhanis YD. 1970 Studies on the bioluminescence of Renilla reniformis. VII. Conversion of luciferin into luciferyl sulfate by luciferin sulfokinase. Biochemistry 9, 1184–1189.

44. Oakley TH. 2005 Myodocopa (Crustacea: Ostracoda) as models for evolutionary studies of light and vision: multiple origins of bioluminescence and extreme sexual dimorphism. Hydrobiologia 538, 179–192.

45. Abe K, Ono T, Yamada K, Yamamura N, Ikuta K. 2000 Multifunctions of the upper lip and a ventral reflecting organ in a bioluminescent ostracod Vargula hilgendorfii (Müller, 1890). Hydrobiologia

46. Losos JB. 2011 Convergence, adaptation, and constraint. Evolution 65, 1827–1840.

47. Wake DB. 1991 Homoplasy: The Result of Natural Selection, or Evidence of Design Limitations? Am. Nat. 138, 543–567.

