## Supplemental Material for "Similar enzymatic functions in distinct bioluminescence systems: Evolutionary recruitment of sulfotransferases in ostracod light organs"

### Supplemental Figures

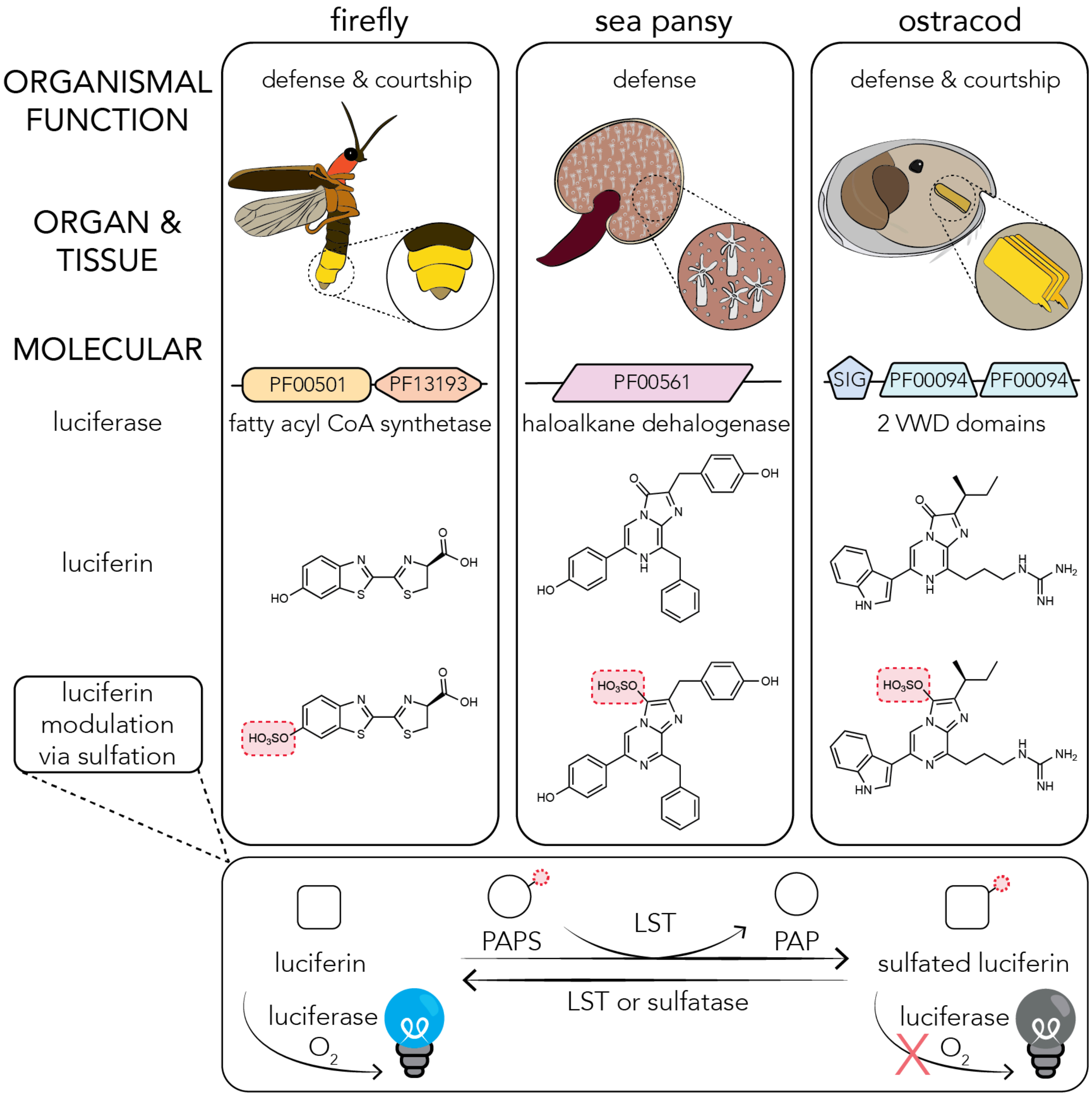

**Figure S1. Convergently evolved bioluminescence systems of fireflies, sea pansies, and ostracods may vary in the extent of convergence across and within levels of biological organization.**

These taxa use bioluminescence as a defense mechanism but produce light by using structurally different organs and tissues, non-homologous luciferases (PF00501 = AMP-dependent synthetase/ligase, PF13193 = AMP-binding enzyme C-terminal domain, PF00561 = alpha/beta hydrolase fold, SIG = signal peptide, PF00094 = von Willebrand factor type D domain), and different luciferins. Despite these differences, they use sulfation as a common mechanism to modulate the chemical state of their luciferins. Luciferin sulfation is catalyzed by a luciferin sulfotransferase (LST), which transfers a sulfo group from the donor 3′-phosphoadenosine-5′-phosphosulfate (PAPS) to luciferin and produces 3’-phosphoadenosine-5’-phosphate (PAP) and sulfated luciferin, which cannot be oxidized by their respective luciferase to emit light. The sulfo group may be removed, either by a LST or sulfatase, to produce the active form of luciferin, which is oxidized by luciferase to produce light.

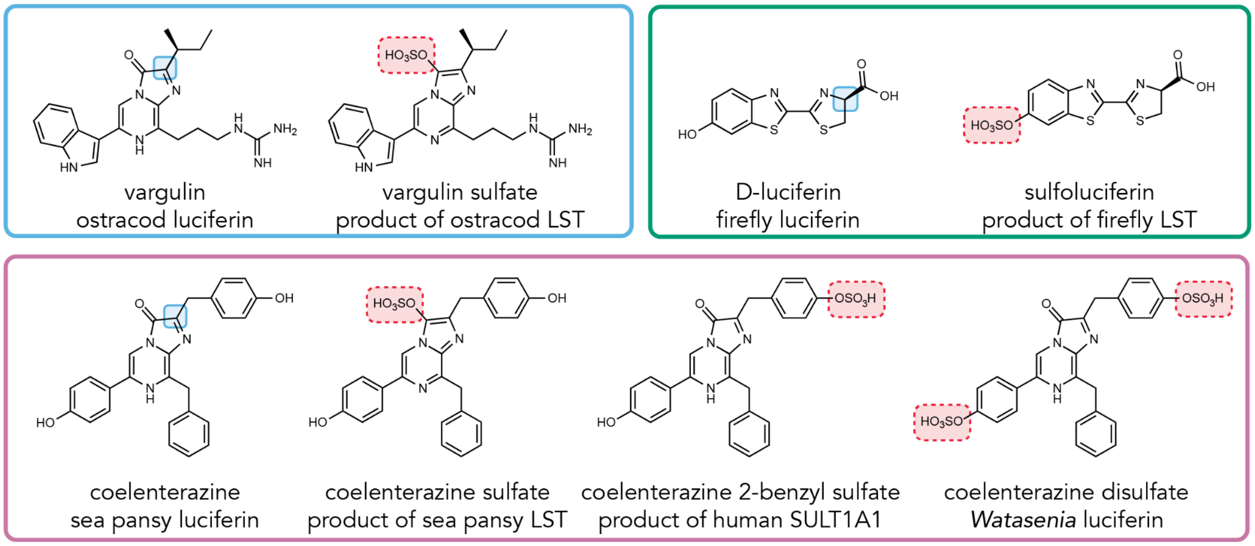

**Figure S2. Chemical structures of luciferins from ostracods (vargulin), sea pansies (coelenterazine), and fireflies (D-luciferin).**

Sites of luciferin modification by oxidation and sulfation are boxed in blue and red, respectively. The sites of sulfation in coelenterazine by sea pansy luciferin sulfotransferase (LST) and human SULT1A1 differ. The products of ostracod, firefly, and sea pansy LSTs are unable to react with their respective luciferases. However, in the firefly squid *Watasenia*, coelenterazine disulfate is used in its bioluminescence reaction.

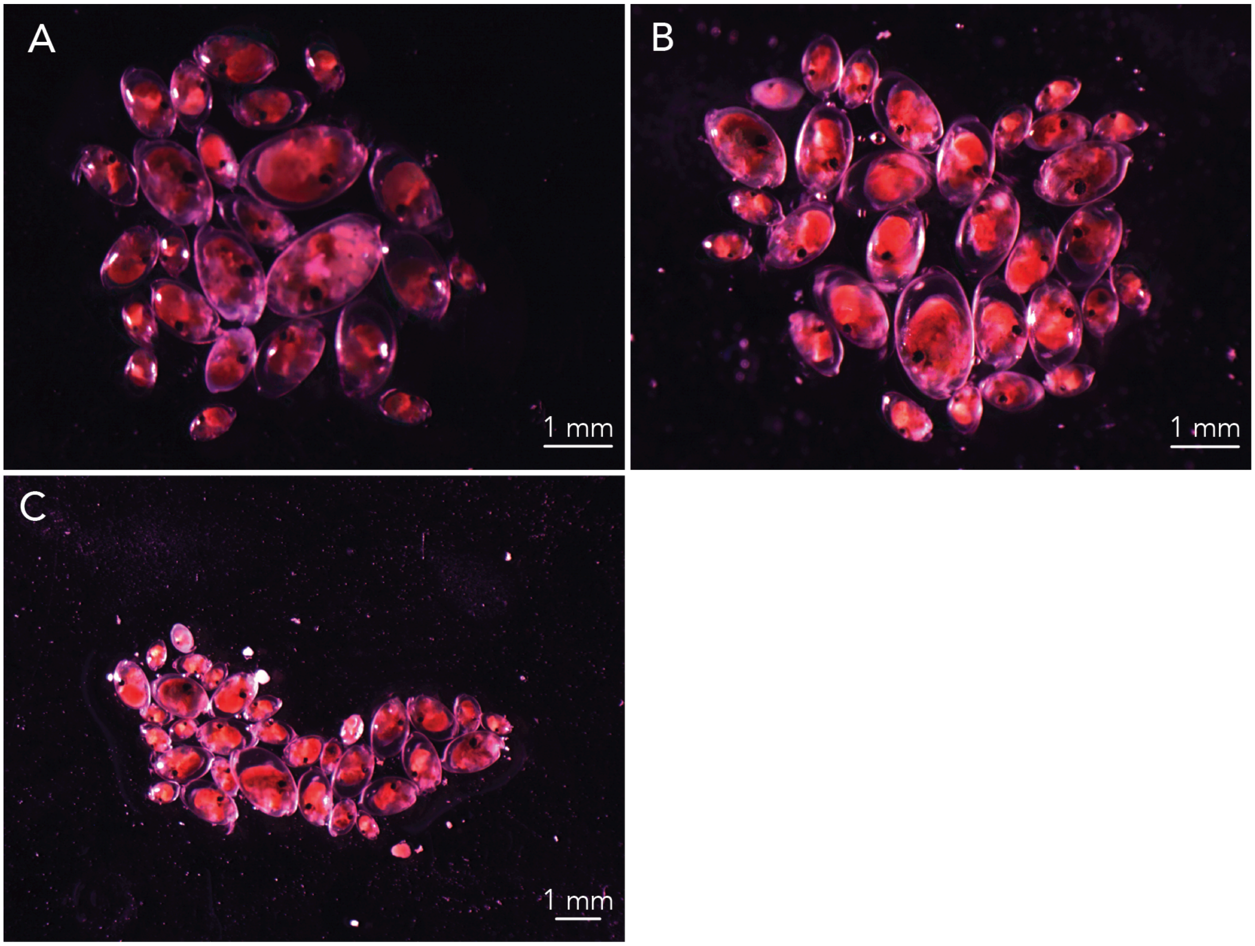

**Figure S3. *Vargula tsujii* specimens sequenced via Iso-Seq to produce the reference transcriptome.**

(A) Image “vt3”, 1.25x magnification. (B) Image “vt4”, 1.25x magnification. (C) Image “vt5”, 0.8x magnification. The names of each image are referred to in Supplemental Table 1, which lists the measurements and inferred instar stage of individuals in each photo.

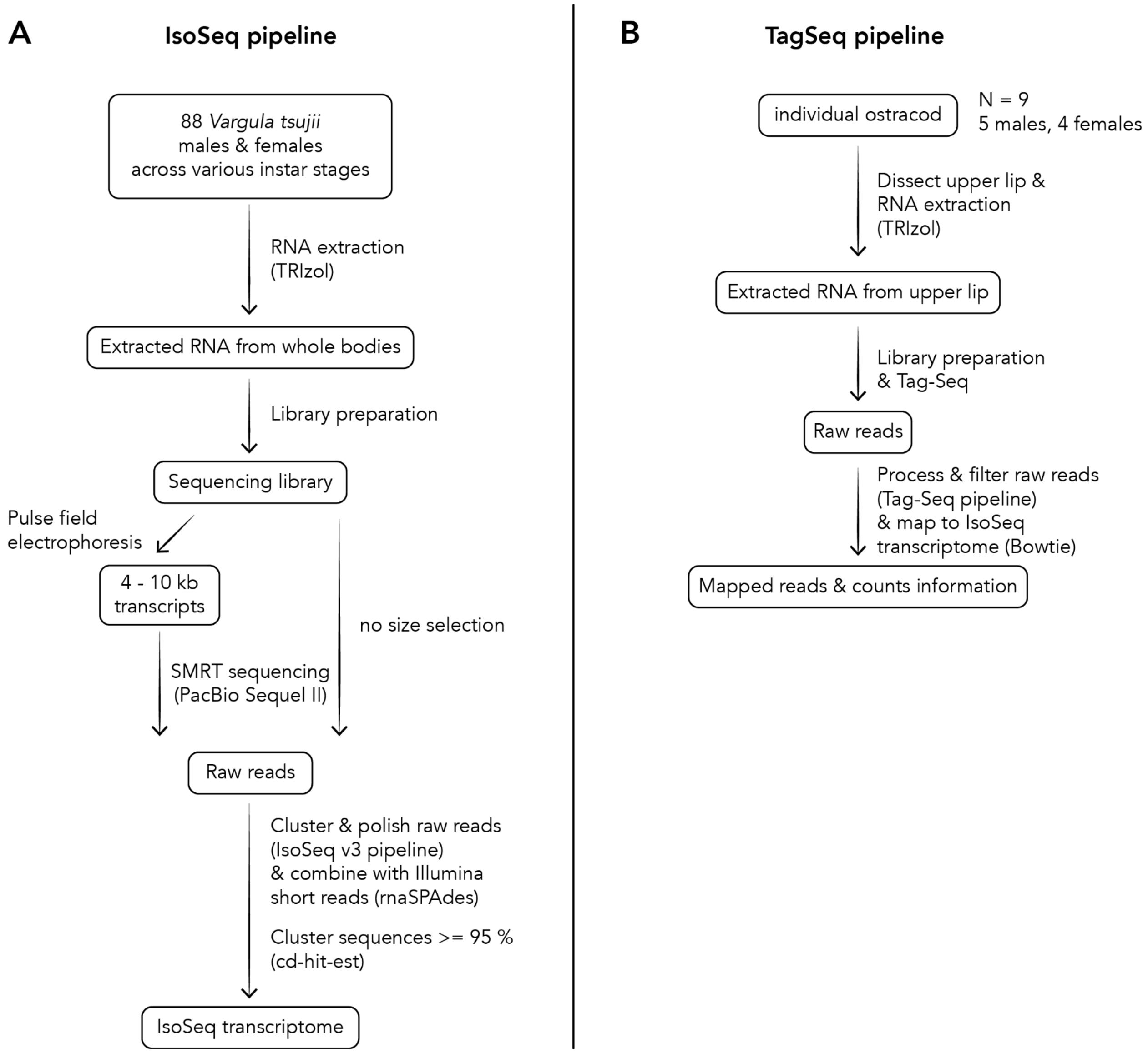

**Figure S4. Flowchart of RNA sequencing and bioinformatics pipelines.**

(A) We produced a high quality reference transcriptome by extracting and sequencing RNA from the whole bodies of 88 individuals using Single Molecular Real-Time (SMRT) Sequencing. (B) To characterize the expression of genes in the light-producing organ — the upper lip — of ostracods, we extracted and sequenced RNA from the upper lips of ostracods (N = 9) using Tag-Seq.

```
>ST1
MGSSHHHHHHGSGLVPRGSASMSDSEVNQEAKPEVKPEVKPETHINLKVSDGSSEIFFKIKKTTPLRRLMEAFAKRQGK
EMDSLRFLYDGIRIQADQTPEDLDMEDNDIIEAHREQIGGAAYPFTIEDYSSDLDPSPKVILQPGNWVMPNGFRPVAEK
IYNMEVREDDIWLLSWPKTGTTWISEMIWTLCHDSDTELAKTIPHFKRWAFPENDSMWADFSDYLRMQLSDGVPFPKEV
VHQLESDSFEDADKMEGPRFVRSHLPICMLPPNLLDTCKCIYMLRNPMDTCVSFYHFMKTVADVEIDIGMYINQFVADE
AYFSPLWAHVKEAWQKRSHPNMMFLYYEELQKDLPASIRKVSKFLNRPVTEEQVEKLTTHLHIDNMRQNPMINMDEMKA
TGRLRPDAMFLRKGVVGDWKNNLTPKMEMQLINWWNNNMRDCDIRFPGIQLKSRF*
>ST2
MGSSHHHHHHGSGLVPRGSASMSDSEVNQEAKPEVKPEVKPETHINLKVSDGSSEIFFKIKKTTPLRRLMEAFAKRQGK
EMDSLRFLYDGIRIQADQTPEDLDMEDNDIIEAHREQIGGKKVPFTYEDCKVKDDDGEYVRVKPGNFLLPATIRDYGEE
IYNMELREGDVILLSWPKTGTHWCSELIWNICNDMNIEEAKSKSMLDRWRFIDVPSDKKKHVCALELTDKPMGRIEKLL
DVEGPRYIISHLPLSLFPPDTLDKCKVVYVTRNPMDTIVSLYRFLELRPDENTDFPTFVNNFMHDEVVFSPYWEHVGEA
WKLKEHSNLFFLTFEEMKKDLRATIKNVAKFFNKEFSIEQIDTLVDHLSFENMQSNKSVNAEYLNKTGKIKQGGQFIRK
GAVGDFKNSFTPAMQSQMKNWIQTNNRGCDIKFQGVAM*
>ST3
MRGSHHHHHHMTTFPYKFEEYQGGVADAIKLNPGGWVMLSTYQEDAEKLYNFGARPDDIWLVGHPKTGTTWSSELLWTV
LHDADVEGARTIPQMDRICYPEYDFIIVKRDLADMVQECKKLGKEVPEKLLKESTKSHYQMAVEHPSPRLLRTGMCLEL
LSPKLLDTCKVVYVTRNPLDTCVSFYHFINSMEATKEHVDFQDFFQLFLADLVPFGPFWKHVKYAWEKRHHPNMLFLFY
EDMKADLPGVIKKVADFVGKSLTEEQIERVAQYLSFDEMKKNKMVNLDSEREMGVWKKDSDFVRKGIVGDYKNHMTPEM
IEQLRKWWKEHMEDCDIKFPGFDEL*
>ST4
MRGSHHHHHHMRIQTRCVISERNMNEIPFTLEEFTSEDDPRRKVRVQPINYVLFAECKELLTRIYNLKFRPGDIVLITT
PKSGTHWLIEMLWTMINNVDTEAAKKDVHWERAPMLEINHIDIDVVKELKEKRQNGEKLMPMEDFWVDYDSLKKFEALP
SPRVCFTHLPIPMLPPNMLEVCKVIYLTRNPADVICSYHQFMHDRCHLHVDFDTFFNMFLDGTVLFTPYWSSVKEGWKI
RNHPNALFLLYEEMAKDLRGCIEKISHHIQCPLSEEQIDKLQDHLSFENMKKNPAINMTPLAGTAFMKKGGNVVRKGKI
GDSKTTLSEEQNKRLKDWCEANREECDIKFPGSDF*
>ST5
MGSSHHHHHHGSGLVPRGSASMSDSEVNQEAKPEVKPEVKPETHINLKVSDGSSEIFFKIKKTTPLRRLMEAFAKRQGK
EMDSLRFLYDGIRIQADQTPEDLDMEDNDIIEAHREQIGGADLPFTLEEFEVAEESRLKVRVQPGNFTLYAGYKQLASR
IYNIKFRSGDVVLMSTPKSGTHWAAEMLWNIINQVDIEAAKKDLLFRRVPMLEISHIDAEKTEELKEKKAAGENLLPIE
ETVIFNDSIETFEKLPSPRICYTHLPFILLPPNILEVCKIVYITRNPADVIVSYHPFMAAGPNFDVNFENFFKMFLDGT
ALFTPYWSSVKEAWKRKDHPNMLFLLYEEMKQDLRGCIKNVSKHLQRPLTGEQVDRLEDHLSFESMKSNSSVNMKHLAG
SAFFSKDGQLCRKGQIGDSLNTLSDDQKKQMKEWCDANRGDCNITFPGLDQ*
>C.noctiluca_luciferase
MKTLILAVALVYCATVHCQDCPYEPDPPNTVPTSCEAKEGECIDSSCGTCTRDILSDGLCENKPGKTCCRMCQYVIECR
VEAAGWFRTFYGKRFQFQEPGTYVLGQGTKGGDWKVSITLENLDGTKGAVLTKTRLEVAGDIIDIAQATENPITVNGGA
DPIIANPYTIGEVTIAVVEMPGFNITVIEFFKLIVIDILGGRSVRIAPDTANKGMISGLCGDLKMMEDTDFTSDPEQLA
IQPKINQEFDGCPLYGNPDDVAYCKGLLEPYKDSCRNPINFYYYTISCAFARCMGGDERASHVLLDYRETCAAPETRGT
CVLSGHTFYDTFDKARYQFQGPCKEILMAADCFWNTWDVKVSHRNVDSYTEVEKVRIRKQSTVVELIVDGKQILVGGEA
VSVPYSSQNTSIYWQDGDILTTAILPEALVVKFNFKQLLVVHIRDPFDGKTCGICGNYNQDFSDDSFDAEGACDLTPNP
PGCTEEQKPEAERLCNSLFAGQSDLDQKCNVCHKPDRVERCMYEYCLRGQQGFCDHAWEFKKECYIKHGDTLEVPDECK
GPDYKDHDGDYKDHDIDYKDDDDKGPHHHHHH*
```

**Figure S5. Gene sequences of recombinant proteins produced in this study.**

Proteins were tagged with a hexa-histidine tag (shown in orange) to facilitate purification by immobilized metal affinity chromatography (IMAC) using Ni-NTA resin. To improve solubility, ST1, ST2, and ST5 were engineered to incorporate a SUMO tag (shown in blue) at the N terminus. Luciferase was further tagged with a 3 x FLAG epitope (shown in green) and hexa-histidine tag at the C terminus.

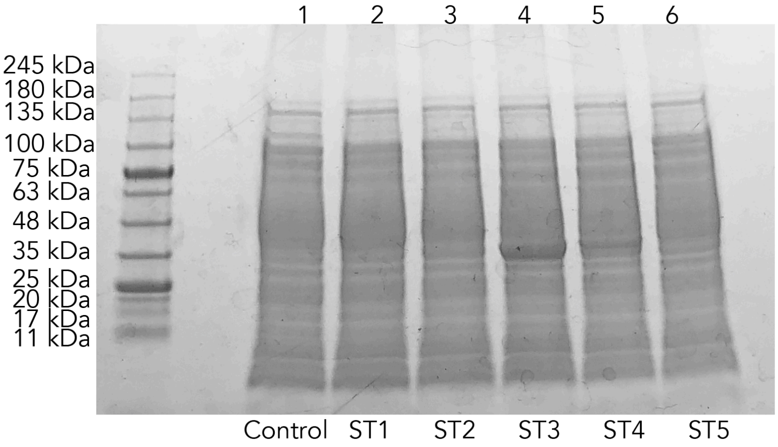

**Figure S6. SDS-PAGE of crude lysates prepared from *E. coli* BL21 cells induced to express ST1-5.**

Lane 1: cells transformed with control pQE80 vector lacking any ST, lane 2: ST1, lane 3: ST2, lane 4: ST3, lane 5: ST4, and lane 6: ST5 expressing *E. coli* cell lysates. SUMO-tags were not included for any ST construct. Bands at the expected size were detected only for ST3 (40.11 kDa) and ST4 (41.29 kDa).

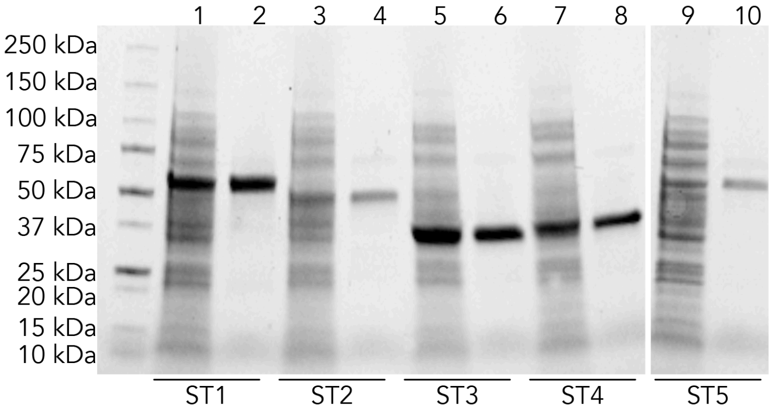

**Figure S7. Polyacrylamide gel electrophoresis of purified sulfotransferase candidates.**

Lanes 1, 3, 5, 7, and 9 contain (respectively) crude lysates prepared from ST1, ST2, ST3, ST4, and ST5 expressing *E. coli* cells. Lanes 2, 4, 6, 8, and 10 contain purified protein fractions of ST1, ST2, ST3, ST4, and ST5 prepared by IMAC with Ni-NTA resin. ST1, ST2, and ST5 are tagged with a SUMO peptide to increase solubility.

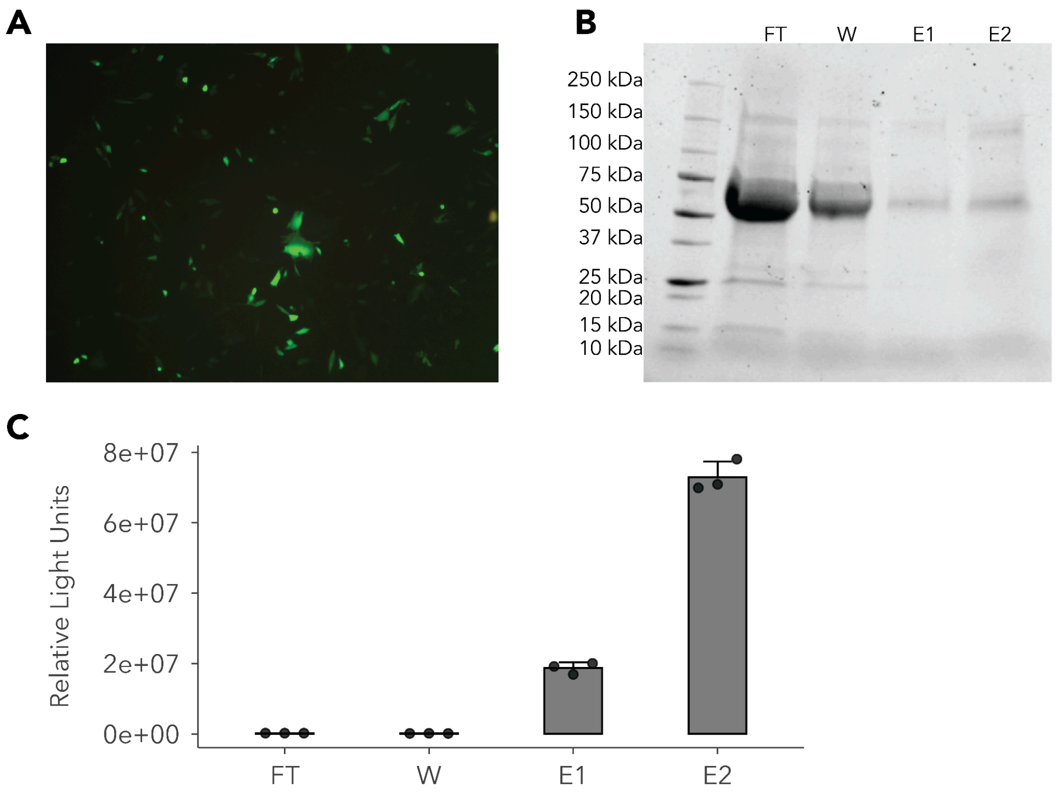

**Figure S8. Expression and purification of ostracod luciferase.**

(A) We co-transfected Chinese hamster ovary cells with the modified ostracod luciferase (from *Cypridina noctiluca*) vector and a GFP marker plasmid. We monitored expression of GFP in CHO cells to confirm transfection. Cells were excited with blue light and visualized using a 10X objective lens (Olympus CKX-41). 48 hours post transfection, we harvested the media, which contains the secreted luciferase, and purified the ostracod luciferase by performing IMAC using Ni-NTA agarose resin. (B) SDS-PAGE of each purification step. Lane 1: flow-through (FT); lane 2: washout fraction (W) from treatment with 10 mM imidazole containing buffer; lanes 3 and 4: elution fractions (E1 and E2) following treatment with 500 mM imidazole. (C) Presence of luciferase was verified by recording luminescence produced by incubating samples from each purification step with luciferin (N = 3).

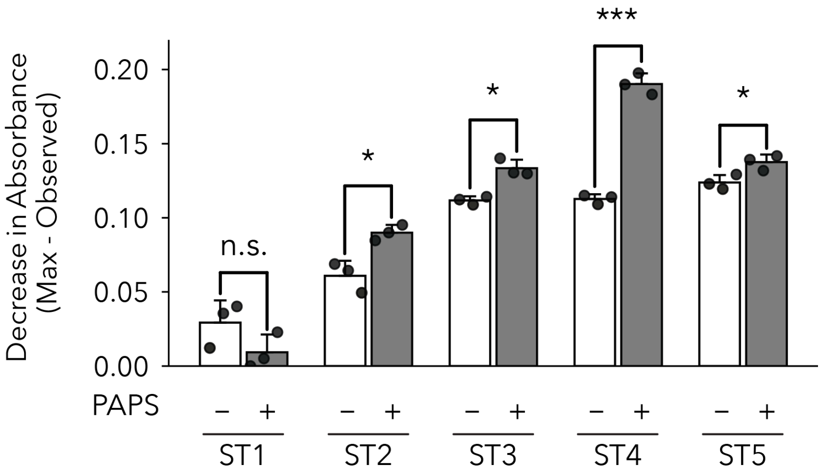

**Figure S9. Four candidates, especially ST4, sulfate p-nitrophenol under the experimental conditions tested.**

Sulfation of p-nitrophenol, a common substrate for many sulfotransferases, was assayed by measuring change in absorbance at 405 nm following a 15 hour incubation with purified sulfotransferase proteins in the absence and presence of PAPS. Control reactions, in white, contained 100 μL of 4 μM purified sulfotransferase and 100 μM p-nitrophenol but omitted PAPS to prevent sulfation activity. Experimental reactions, in gray, contained 100 μL of 4 μM purified sulfotransferase, 100 μM p-nitrophenol, and 500 μM PAPS. Four candidates — ST2, ST3, ST4, and ST5 — catalyze statistically significant p-nitrophenol sulfation under the conditions tested, with ST4 by far the highest effect size. Error bars represent the standard deviation (N = 3). * denotes $P < 0.05$, ** denotes $P < 0.01$, *** denotes $P < 0.001$, and n.s. denotes non-significance ($P \geq 0.05$). P values were calculated by performing Welch's t-tests between STs incubated with and without PAPS.

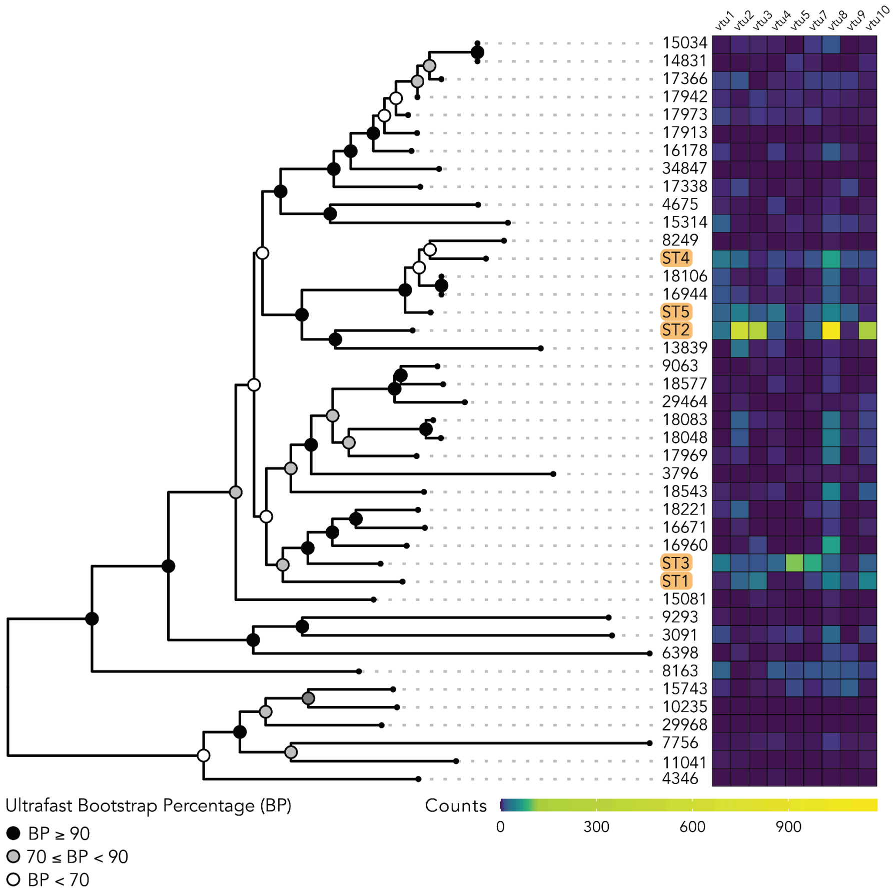

**Figure S10. Phylogeny and gene expression of sulfotransferases from 9 individuals of *Vargula tsujii*.**

A maximum-likelihood phylogeny was inferred using candidate sulfotransferases identified from the Iso-Seq reference transcriptome. Raw reads from Tag-seq were filtered and mapped to the reference transcriptome. The color bar represents read counts in each ostracod sample. Ultrafast bootstrap values are represented by colored circles at each node.

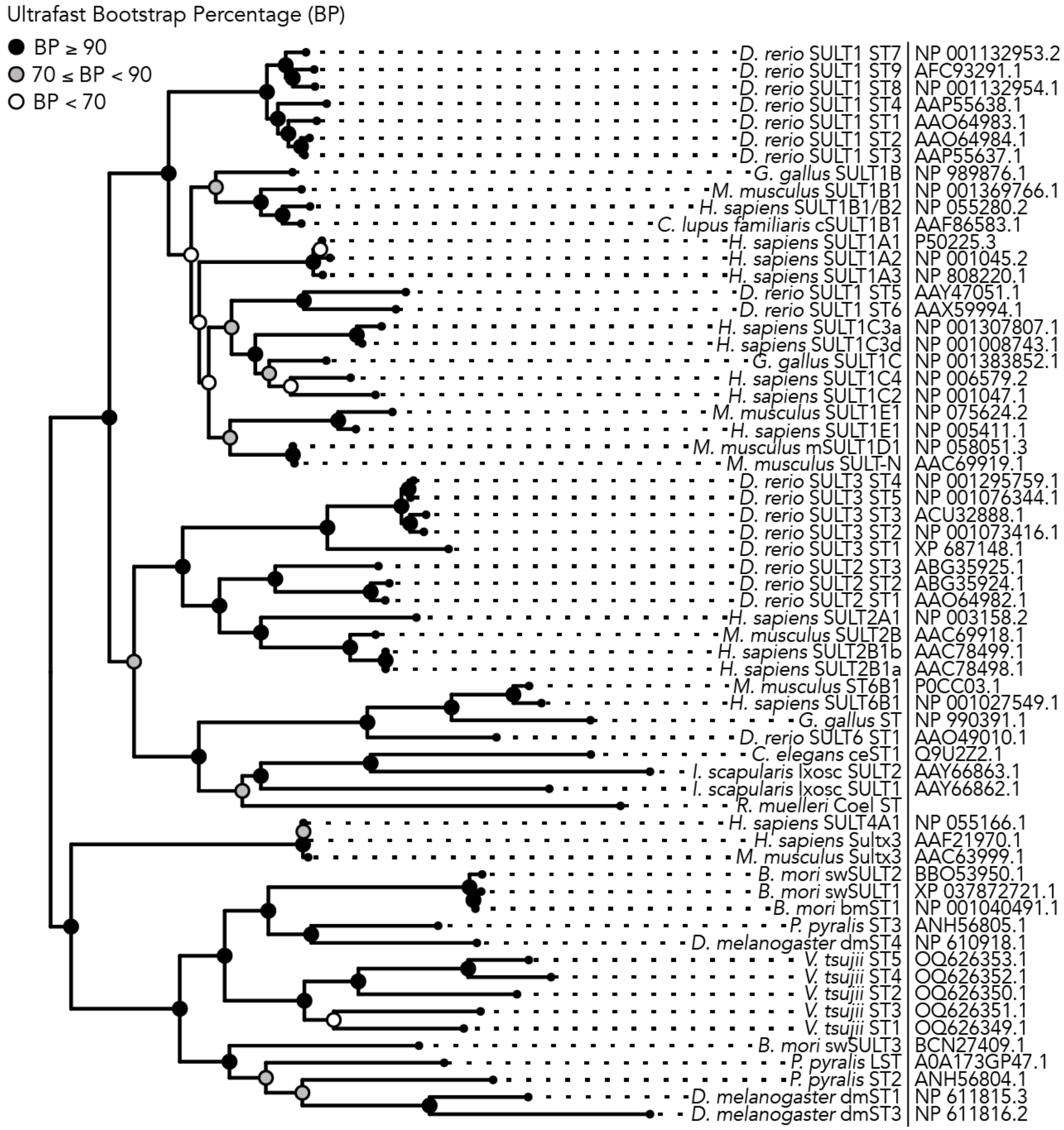

**Figure S11. Phylogeny of functionally tested sulfotransferases, inferred from a trimmed alignment.**

Maximum-likelihood phylogeny using a trimmed alignment of functionally tested sulfotransferases from the literature (as of March 2023). Names of organisms and the sulfotransferases are at the tips, and accession numbers are listed on the right.

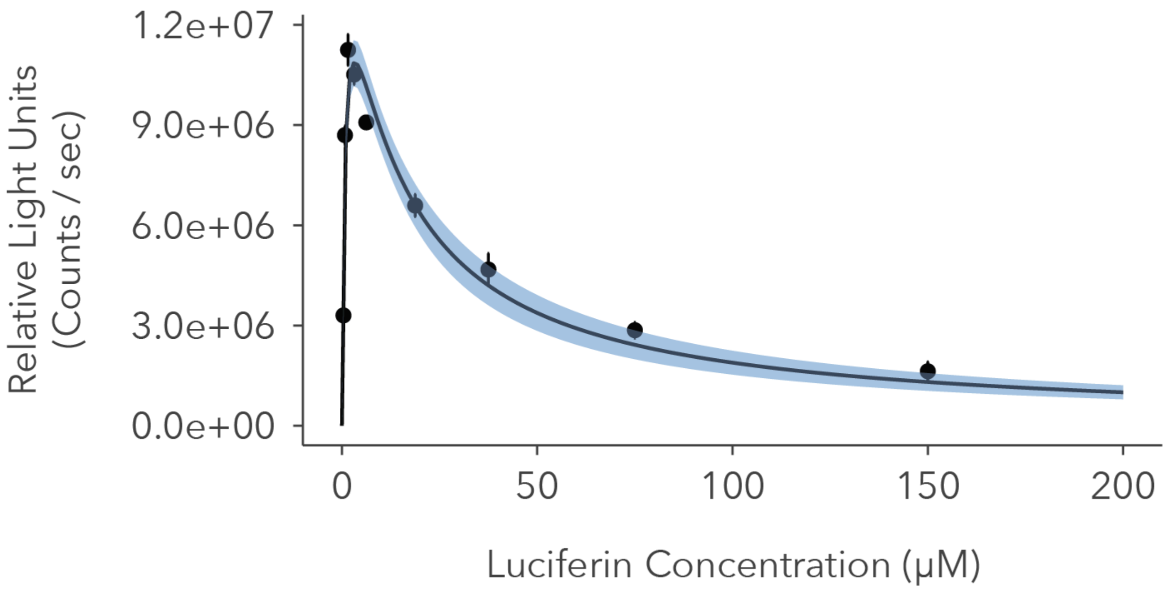

**Figure S12. Ostracod luciferase exhibits substrate inhibition.**

Bioluminescence produced by ostracod luciferase increases as substrate concentration increases — and peaks at a final substrate concentration of 1.5625 μM — but decreases at final substrate concentrations equal to or greater than 3.125 μM, suggesting ostracod luciferase exhibits substrate inhibition. We used a multi-channel pipette to manually add 65 μL of purified ostracod luciferase to 10 μL of luciferin, which resulting in a final concentration of 250 nM luciferase and varying final concentrations — 0.39 μM, 0.78125 μM, 1.5625 μM, 3.125 μM, 6.25 μM, 18.75 μM, 37.5 μM, 75 μM, and 150 μM — of luciferin. We incubated the reactions at room temperature for 12 minutes to stabilize the reading and quantified the luminescence using a microplate reader (Tecan Spark) with an integration time of 2 seconds. Data were fitted using a Michaelis-Menten model modified for substrate inhibition — $v = V_{max}$ [S] / $K_m$ + [S] (1 + [S] / $K_i$) — where v = velocity (relative light units produced per second), $V_{max}$ = maximum velocity, S = substrate concentration, $K_m$ = Michaelis-Menten constant, and $K_i$ = the dissociation constant for the inactive complex formed when two substrate molecules bind to an enzyme (equation 5.43 in [1]). Shaded areas represent 90 % confidence intervals. Estimated $K_m$ = 0.8288 μM, $V_{max}$ = 1.634e7, and $K_i$ = 13.09 μM.

```
                                                                                                                              5′ PSB
            1
SULT_1A1    ---------- ---------- -MELIQDTSR P--------- ---------- ----P--LEY VK----GVPL IKYFAEALGP LQSFQARPDD LLISTYPKSG TTWVSQILDM
Vtsu_ST1    ---------- ---MAAYP-F TIEDYSSDLD PSPKVI---- ---------- --LQP--GNW V--------M PNGFRPVAEK IYNMEVREDD IWLLSWPKTG TTWISEMIWT
Vtsu_ST2    ---------- ---MKKVP-F TYEDCKVKDD DGEYVR---- ---------- --VKP--GNF L--------L PATIRDYGEE IYNMELREGD VILLSWPKTG THWCSELIWN
Vtsu_ST3    ---------- ---MTTFP-Y KFEEYQGGVA DAIK------ ---------- --LNP--GGW V--------M LSTYQEDAEK LYNFGARPDD IWLVGHPKTG TTWSSELLWT
Vtsu_ST4    MRIQTRCVIS ERNMNEIP-F TLEEFTSEDD PRRKVR---- ---------- --VQP--INY V--------L FAECKELLTR IYNLKFRPGD IVLITTPKSG THWLIEMLWT
Vtsu_ST5    ---------- ---MADLP-F TLEEFEVAEE SRLKVR---- ---------- --VQP--GNF T--------L YAGYKQLASR IYNIKFRSGD VVLMSTPKSG THWAAEMLWN
LST         MFASI----- ---LGKIPQY STISSKCSLK ECRKVKKVEN SDEIGEIIHR DFLDPFCNEY VLIGEEETSL GINYLQFEEE IRTFEIRDSD IIVASYPKAG TTWTQELVWL
Coel_ST     ---------- ---MS----A TKETQSSNEF DFPAIR---- ---------- -------ERI V--------S IVTSEESCKK ALEFVPRSTD IFIVTPPKCG TTWMQQIVHQ

          111
SULT_1A1    IYQGGDLEKC HRAPIFMR-- ---VPFLEF- ---------- ---KAPGI-- ---------- ---PSGMETL KDTPAPRLLK THLPLALLPQ TLLD--QKVK VVYVARNAKD
Vtsu_ST1    LCHDSDTELA KTIPHFKRWA ---FPENDSM WA--DFSDYL RMQLSDGVPF PKEVVHQLE- ---SDSFEDA DKMEGPRFVR SHLPICMLPP NLLD---TCK CIYMLRNPMD
Vtsu_ST2    ICNDMNIEEA KSKSMLDRWR FIDVPSDKKK HV---CALEL TDKPMGRI-- -EKLLDVE-- ---------- ----GPRYII SHLPLSLFPP DTLD---KCK VVYVTRNPMD
Vtsu_ST3    VLHDADVEGA RTIPQMDRIC ---YPEYDFI IVKRDLADMV QECKKLGKEV PEKLLKEST- ---KSHYQMA VEHPSPRLLR TGMCLELLSP KLLD---TCK VVYVTRNPLD
Vtsu_ST4    MINNVDTEAA KKDVHWER-- ---APMLEIN HIDIDVVKEL KEKRQNG--- -EKLMPMEDF WVDYDSLKKF EALPSPRVCF THLPIPMLPP NMLE---VCK VIYLTRNPAD
Vtsu_ST5    IINQVDIEAA KKDLLFRR-- ---VPMLEIS HIDAEKTEEL KEKKAAG--- -ENLLPIEET VIFNDSIETF EKLPSPRICY THLPFILLPP NILE---VCK IVYITRNPAD
LST         IGNDLDFKAA EE--HLDK-- --RFPHFELC TI----VNFA KMTEMLGSTR PEYI------ ---GNSINYL RDLEGTRFIK THLTYNLLPE QILNGNRKPK IIYVMRDPKD
Coel_ST     LQTGGDMD-- -----FDEIH -AEIPWIEMA HD--IGIDLE KDQKV----- ---------- ---------- ----ATRVFK THLPYDPCPK G------ASK YIVVFRNPCS

          221
SULT_1A1    VAVSYYHFYH MAKVHPEPGT WDSF-LEKFM -------VGE VSYGSWYQHV QEWWELSRTH P-VLYLFYED MKENPKREIQ KILEFVGRSL PEETVDFVVQ HTSFKEMKKN
Vtsu_ST1    TCVSFYHFMK TVADVE--ID IGMY-INQFV -------ADE AYFSPLWAHV KEAWQK-RSH PNMMFLYYEE LQKDLPASIR KVSKFLNRPV TEEQVEKLTT HLHIDNMRQN
Vtsu_ST2    TIVSLYRFLE LRPDEN--TD FPTF-VNNFM -------HDE VVFSPYWEHV GEAWKL-KEH SNLFFLTFEE MKKDLRATIK NVAKFFNKEF SIEQIDTLVD HLSFENMQSN
Vtsu_ST3    TCVSFYHFIN SMEATKEHVD FQDF-FQLFL -------ADL VPFGPFWKHV KYAWEK-RHH PNMLFLFYED MKADLPGVIK KVADFVGKSL TEEQIERVAQ YLSFDEMKKN
Vtsu_ST4    VICSYHQFMH DRCHLH--VD FDTF-FNMFL -------DGT VLFTPYWSSV KEGWKI-RNH PNALFLLYEE MAKDLRGCIE KISHHIQCPL SEEQIDKLQD HLSFENMKKN
Vtsu_ST5    VIVSYHPFMA AGPNFD--VN FENF-FKMFL -------DGT ALFTPYWSSV KEAWKR-KDH PNMLFLLYEE MKQDLRGCIK NVSKHLQRPL TGEQVDRLED HLSFESMKSN
LST         VCVSYYHHGR LIQGWR--AD FQNF-SKVFL -------SEK IMFGSYWKHV LGYWEH-RDK PNVLILTYEE MKKDLLSVIR KTAQFLDKKL NENKIPQLLK HLSFESMKNN
Coel_ST     AAYSFYKF-Q LGALIPENLS VKDYVFEIWL QMGNPPDAKQ PWWVSYFSFL KSWLPH-KND PNVLFLFFEE MKENLENVVR AVSAFLGVN- NEESIRAAVE MSSFDFMK--

                                    GXX GXXK
          331
SULT_1A1    PMTNYTTVPQ EFMDHS---- ISPFMRKGMA GDWKTTFTVA QNERFDADYA EKMAGCSLSF RSEL------ --------
Vtsu_ST1    PMINMDEMKA TGRLRP---- DAMFLRKGVV GDWKNNLTPK MEMQLINWWN NNMRDCDIRF PGI-QLKSRF --------
Vtsu_ST2    KSVNAEYLNK TGKIKQ---- GGQFIRKGAV GDFKNSFTPA MQSQMKNWIQ TNNRGCDIKF QGVAM----- --------
Vtsu_ST3    KMVNLDSERE MGVWKK---- DSDFVRKGIV GDYKNHMTPE MIEQLRKWWK EHMEDCDIKF PGFDEL---- --------
Vtsu_ST4    PAINMTPLAG TAFMKK---- GGNVVRKGKI GDSKTTLSEE QNKRLKDWCE ANREECDIKF PGSDF----- --------
Vtsu_ST5    SSVNMKHLAG SAFFSK---- DGQLCRKGQI GDSLNTLSDD QKKQMKEWCD ANRGDCNITF PGLDQ----- --------
LST         RAVNQQDKIE SRMKHKLVPE QGAFMRSGTS QNYKGEMSEE LILKFDEWTK KSISGSSFVT EPIADLYLNT SQKSVPKM
Coel_ST     --TNADKFDP KPIANKPLPK PMIVIRTGSS TEAMQVLTDE IKEAIQKRWE NEITE-SIGF KTYDELRYAI REECPLKY
```

**Figure S13. Multiple sequence alignment of luciferin sulfotransferases.**

Sulfotransferase domains (Pfam ID: PF00685), identified by hmmscan, are highlighted in each sequence (human SULT1A1 = gray, *V. tsujii* STs = blue, firefly LST = green, *Renilla* ST = purple). The annotated regions above the alignment correspond to the 5’ phosphate-sulfate binding (PSB) loop (sequence TYPKSGT [2,3]) and the GXXGXXK motif [4], two important sites for PAPS binding, in human SULT1A1.

### Supplemental Tables

| Image | Animal | Length (mm) | Inferred stage |
|---|---|---|---|
| vt3 | 1 | 1.25 | A-II |
| vt3 | 2 | 0.78 | A-V |
| vt3 | 3 | 1.17 | A-II |
| vt3 | 4 | 1.076 | A-III |
| vt3 | 5 | 1.173 | A-II |
| vt3 | 6 | 1.141 | A-III |
| vt3 | 7 | 1.582 | A-I/A Male |
| vt3 | 8 | 0.988 | A-III |
| vt3 | 9 | 1.972 | A Female |
| vt3 | 10 | 1.462 | A-I |
| vt3 | 11 | 1.2 | A-II |
| vt3 | 12 | 1.144 | A-III |
| vt3 | 13 | 0.76 | A-V |
| vt3 | 14 | 1.609 | A-I/A Male |
| vt3 | 15 | 2.018 | A Female |
| vt3 | 16 | 1.512 | A-I |
| vt3 | 17 | 0.68 | A-V |
| vt3 | 18 | 0.819 | A-IV |
| vt3 | 19 | 1.244 | A-II |
| vt3 | 20 | 0.769 | A-V |
| vt3 | 21 | 1.064 | A-III |
| vt3 | 22 | 1.403 | A-I |
| vt3 | 23 | 1.504 | A-I |
| vt3 | 24 | 0.738 | A-V |
| vt3 | 25 | 0.792 | A-IV |
| vt4 | 1 | 0.806 | A-IV |
| vt4 | 2 | 0.954 | A-IV |
| vt4 | 3 | 0.925 | A-IV |
| vt4 | 4 | 1.502 | A-I |
| vt4 | 5 | 1.398 | A-I |
| vt4 | 6 | 1.35 | A-I |
| vt4 | 7 | 1.405 | A-I |
| vt4 | 8 | 1.352 | A-I |
| vt4 | 9 | 0.701 | A-V |
| vt4 | 10 | 0.761 | A-V |
| vt4 | 11 | 0.997 | A-III |
| vt4 | 12 | 0.815 | A-IV |
| vt4 | 13 | 1.556 | A-I/A Male |
| vt4 | 14 | 0.903 | A-IV |
| vt4 | 15 | 0.727 | A-V |
| vt4 | 16 | 1.257 | A-II |
| vt4 | 17 | 1.434 | A-I |
| vt4 | 18 | 1.457 | A-I |
| vt4 | 19 | 1.073 | A-III |
| vt4 | 20 | 1.445 | A-I |
| vt4 | 21 | 1.082 | A-III |
| vt4 | 22 | 1.482 | A-I |
| vt4 | 23 | 1.985 | A Female |
| vt4 | 24 | 1.416 | A-I |

| | | | |
|---|---|---|---|
| vt4 | 25 | 1.365 | A-I |
| vt4 | 26 | 0.915 | A-IV |
| vt4 | 27 | 0.802 | A-IV |
| vt4 | 28 | 0.943 | A-IV |
| vt4 | 29 | 0.817 | A-IV |
| vt4 | 30 | 0.957 | A-IV |
| vt4 | 31 | 0.781 | A-V |
| vt5 | 1 | 1.298 | A-II |
| vt5 | 2 | 0.692 | A-V |
| vt5 | 3 | 0.729 | A-V |
| vt5 | 4 | 0.743 | A-V |
| vt5 | 5 | 1.43 | A-I |
| vt5 | 6 | 0.926 | A-IV |
| vt5 | 7 | 0.77 | A-V |
| vt5 | 8 | 0.85 | A-IV |
| vt5 | 9 | 0.712 | A-V |
| vt5 | 10 | 1.473 | A-I |
| vt5 | 11 | 0.835 | A-IV |
| vt5 | 12 | 1.415 | A-I |
| vt5 | 13 | 1.449 | A-I |
| vt5 | 14 | 0.94 | A-IV |
| vt5 | 15 | 1.393 | A-I |
| vt5 | 16 | 0.994 | A-III |
| vt5 | 17 | 1.999 | A Female |
| vt5 | 18 | 1.09 | A-III |
| vt5 | 19 | 1.099 | A-III |
| vt5 | 20 | 1.443 | A-I |
| vt5 | 21 | 0.869 | A-IV |
| vt5 | 22 | 0.723 | A-V |
| vt5 | 23 | 1.414 | A-I |
| vt5 | 24 | 0.952 | A-IV |
| vt5 | 25 | 0.722 | A-V |
| vt5 | 26 | 1.455 | A-I |
| vt5 | 27 | 1.501 | A-I |
| vt5 | 28 | 1.425 | A-I |
| vt5 | 29 | 1.445 | A-I |
| vt5 | 30 | 0.91 | A-IV |
| vt5 | 31 | 1.581 | A-I/A Male |
| vt5 | 32 | 0.817 | A-IV |

**Table S1. Carapace measurements of *Vargula tsujii* specimens sequenced via Iso-Seq.**

We sexed animals and for each specimen, measured the length of rostrum to keel. We inferred developmental stages based on the classification from [5].

| Sample | Stage | Tissue | Raw Read Counts | Filtered Read Counts | % Raw Reads Kept | Mapped Reads | % Filtered Reads Mapped |
|---|---|---|---|---|---|---|---|
| vtu1 | Adult Male | upper lip | 3,784,061 | 1,210,393 | 32.0% | 989,023 | 81.7% |
| vtu2 | Adult Female | upper lip | 2,508,701 | 607,251 | 24.2% | 450,858 | 74.2% |
| vtu3 | Adult Male | upper lip | 3,286,003 | 1,166,599 | 35.5% | 908,485 | 77.9% |
| vtu4 | Adult Female | upper lip | 3,785,756 | 1,094,550 | 28.9% | 916,547 | 83.7% |
| vtu5 | Adult Male | upper lip | 2,845,340 | 1,021,284 | 35.9% | 740,402 | 72.5% |
| vtu7 | Adult Male | upper lip | 5,725,267 | 2,098,711 | 36.7% | 1,506,546 | 71.8% |
| vtu8 | Adult Female | upper lip | 2,725,196 | 653,138 | 24.0% | 504,336 | 77.2% |
| vtu9 | Adult Male | upper lip | 3,486,104 | 1,211,298 | 34.7% | 933,095 | 77.0% |
| vtu10 | Adult Female | upper lip | 4,071,147 | 1,197,806 | 29.4% | 909,492 | 75.9% |

**Table S2. Tag-Seq sequencing, filtration, and mapping statistics.**

| Protein | Forward Gibson primer 5’-3’ | Reverse Gibson primer 5’-3’ |
|---|---|---|
| ST1 | AGCTGAAGAGCCGTTTCTGAAGA<br>CAAGCTTAGGTATTTATTCGGCG | ATGGTAAACGGATACGCAGCACC<br>ACCAATCTGTTCTCTGTGAG |
| ST2 | TCCAAGGGGTTGCTATGTAAAGA<br>CAAGCTTAGGTATTTATTCGGCG | TAAGTAAATGGCACCTTTTTACC<br>ACCAATCTGTTCTCTGTGAG |
| ST3 | CAGGTTTCGATGAGCTTTAAAAG<br>CTTAATTAGCTGAGCTTGGACTC | TTGTAGGGGAAAGTTGTCATGTG<br>ATGGTGATGGTGATGCGAT |
| ST4 | TCCCCGGCTCCGATTTCTGAAAG<br>CTTAATTAGCTGAGCTTGGACTC | CAACGCGTCTGAATACGCATGTG<br>ATGGTGATGGTGATGCGAT |
| ST5 | TTCCCGGTTTGGACCAGTGAAGA<br>CAAGCTTAGGTATTTATTCGGCG | AGCGTAAACGGAAGATCGGCACC<br>ACCAATCTGTTCTCTGTGAG |
| Ostracod luciferase | GATTACAAGGATGACGATGACAA<br>GGGCCCTCACCACCACCATCACC<br>ACTGAGCGGCCGCAAAATCA | GATGTCATGATCTTTATAATCAC<br>CGTCATGGTCTTTGTAGTCAGGG<br>CCCTTGCACTCGTCGGGCA |

**Table S3. Oligonucleotide primers used in this work to clone the corresponding protein constructs (indicated in the left column) by Gibson assembly.**

| Protein | Molecular weight with purification/solubility tags (kDa) | Molecular weight without purification/solubility tags (kDa) |
|---|---|---|
| ST1 | 52.47 | 39.2 |
| ST2 | 50.07 | 36.8 |
| ST3 | 40.11 | 38.86 |
| ST4 | 41.29 | 40.03 |
| ST5 | 50.94 | 37.67 |
| Ostracod luciferase | 65.32 (with signal peptide)<br>63.39 (no signal peptide) | 61.48 (with signal peptide)<br>59.54 (no signal peptide) |

**Table S4. Estimated molecular weights of recombinant proteins produced in this work.**

| Protein | Purity |
|---|---|
| ST1 | 98.76 ± 1.64 % |
| ST2 | 95.94 ± 1.93 % |
| ST3 | 98.61 ± 0.26 % |
| ST4 | 98.11 ± 0.15 % |
| ST5 | 93.01 ± 1.48 % |

**Table S5. Percentage purity of *V. tsujii* sulfotransferase candidates expressed in this work, as assessed by densitometry analysis of protein bands on a SDS-PAGE gel.**

Each candidate was recombinantly expressed and purified twice. Densitometry analysis was performed for each purification and the percent purity was averaged. The values of percent purity reported in this table represent the average percent purity, and the + and - signs represent the maximum and minimum percent purity values, respectively, for each purification.

| Organism | SULT name | Accession | Luciferin | p-nitrophenol | Other |
|---|---|---|---|---|---|
| *Bombyx mori* | bmST1 | NP_001040491.1 | | No [6] | Yes [6] |
| *Bombyx mori* | swSULT1 | XP_037872721.1 | | No [7] | Yes [7] |
| *Bombyx mori* | swSULT2 | BBO53950.1 | | No [7] | Yes [7] |
| *Bombyx mori* | swSULT3 | BCN27409.1 | | No [8] | Yes [8] |
| *Caenorhabditis elegans* | ceST1 | Q9U2Z2.1 | | Yes [9] | Yes [9] |
| *Canis lupus familiaris* | cSULT1B1 | AAF86583.1 | | Yes [10] | Yes [10] |
| *Danio rerio* | SULT1 ST1 | AAO64983.1 | | Yes [11] | Yes [11] |
| *Danio rerio* | SULT1 ST2 | AAO64984.1 | | Yes [11] | Yes [11] |
| *Danio rerio* | SULT1 ST3 | AAP55637.1 | | Yes [11] | Yes [11] |
| *Danio rerio* | SULT1 ST4 | AAP55638.1 | | Yes [11] | Yes [11] |
| *Danio rerio* | SULT1 ST5 | AAY47051.1 | | Yes [12] | Yes [12] |
| *Danio rerio* | SULT1 ST6 | AAX59994.1 | | Yes [13] | Yes [13] |
| *Danio rerio* | SULT1 ST7 | NP_001132953.2 | | No [14] | Yes [14] |
| *Danio rerio* | SULT1 ST8 | NP_001132954.1 | | No [14] | Yes [14] |
| *Danio rerio* | SULT1 ST9 | AFC93291.1 | | | Yes [15] |
| *Danio rerio* | SULT2 ST1 | AAO64982.1 | | No [11] | Yes [11] |
| *Danio rerio* | SULT2 ST2 | ABG35924.1 | | No [16] | Yes [16] |
| *Danio rerio* | SULT2 ST3 | ABG35925.1 | | No [16] | Yes [16] |
| *Danio rerio* | SULT3 ST1 | XP_687148.1 | | No [17] | Yes [17] |
| *Danio rerio* | SULT3 ST2 | NP_001073416.1 | | No [17] | Yes [17] |
| *Danio rerio* | SULT3 ST3 | ACU32888.1 | | No [18] | Yes [18] |
| *Danio rerio* | SULT3 ST4 | NP_001295759.1 | | | Yes [15] |
| *Danio rerio* | SULT3 ST5 | NP_001076344.1 | | | Yes [15] |
| *Danio rerio* | SULT6 ST1 | AAO49010.1 | | Yes [19] | Yes [19] |
| *Drosophila melanogaster* | dmST1 | NP_611815.3 | | Yes [20] | Yes [20] |
| *Drosophila melanogaster* | dmST3 | NP_611816.2 | | Yes [20] | Yes [20] |
| *Drosophila melanogaster* | dmST4 | NP_610918.1 | | Yes [20] | Yes [20] |
| *Gallus gallus* | SULT1C | NP_001383852.1 | | Yes [21] | Yes [21] |
| *Gallus gallus* | SULT1B | NP_989876.1 | | Yes [21] | Yes [21] |

| | | | | | |
|---|---|---|---|---|---|
| *Gallus gallus* | ST | NP_990391.1 | | | Yes[22] |
| *Homo sapiens* | Sultx3 | AAF21970.1 | | Yes [23] | Yes [23] |
| *Homo sapiens* | SULT1A1 | P50225.3 | Yes* [24] | Yes [25] | Yes [25] |
| *Homo sapiens* | SULT1A2 | NP_001045 | | Yes [25] | Yes [25] |
| *Homo sapiens* | SULT1A3 | NP_808220.1 | | Yes [25] | Yes [25] |
| *Homo sapiens* | SULT1B1/B2 | NP_055280 | | Yes [26] | Yes [26] |
| *Homo sapiens* | SULT1C2 | NP_001047.1 | | Yes [25] | Yes [25] |
| *Homo sapiens* | SULT1C3a | NP_001307807.1 | | No [27] | Yes [27] |
| *Homo sapiens* | SULT1C3d | NP_001008743.1 | | Yes [27] | Yes [27] |
| *Homo sapiens* | SULT1C4 | NP_006579 | | Yes [25] | Yes [25] |
| *Homo sapiens* | SULT1E1 | NP_005411 | | Yes [28] | Yes [28] |
| *Homo sapiens* | SULT2A1 | NP_003158 | | | Yes [25] |
| *Homo sapiens* | SULT2B1a | AAC78498.1 | | No [29] | Yes [29] |
| *Homo sapiens* | SULT2B1b | AAC78499.1 | | No [29] | Yes [29] |
| *Homo sapiens* | SULT4A1 | NP_055166 | | No [30] | Yes [30] |
| *Homo sapiens* | SULT6B1 | NP_001027549 | | No [30] | Yes [30] |
| *Ixodes scapularis* | Ixosc SULT 1 | AAY66862.1 | | Yes [31] | Yes [31] |
| *Ixodes scapularis* | Ixosc SULT 2 | AAY66863.1 | | Yes [31] | Yes [31] |
| *Mus musculus* | ST6B1 | P0CC03.1 | | No [32] | Yes [32] |
| *Mus musculus* | Sultx3 | AAC63999.1 | | Yes [23] | Yes [23] |
| *Mus musculus* | SULT2B | AAC69918.1 | | No [33] | Yes [33] |
| *Mus musculus* | SULT-N | AAC69919.1 | | Yes [33] | Yes [33] |
| *Mus musculus* | SULT1B1 | NP_001369766.1 | | Yes [34] | Yes [34] |
| *Mus musculus* | SULT1E1 | NP_075624.2 | Yes [35] | No [36] | Yes [36] |
| *Mus musculus* | mSULT1D1 | NP_058051.3 | Yes [35] | Yes [37] | Yes [37] |
| *Photinus pyralis* | Ppyr LST | A0A173GP47.1 | Yes [38] | No [38] | |
| *Photinus pyralis* | PpyrST2 | ANH56804.1 | No [38] | Yes [38] | |
| *Photinus pyralis* | PpyrST3 | ANH56805.1 | No [38] | Yes [38] | |
| *Renilla muelleri* | Coel ST | | Yes [35] | | |
| *Vargula tsujii* | VtST1 | OQ626349 | No | Undetected | |

| | | | | |
|---|---|---|---|---|
| *Vargula tsujii* | VtST2 | OQ626350 | No | Small |
| *Vargula tsujii* | VtST3 | OQ626351 | Yes** | Small |
| *Vargula tsujii* | VtST4 | OQ626352 | No | Large |
| *Vargula tsujii* | VtST5 | OQ626353 | Yes** | Small |

**Table S6. Functionally characterized sulfotransferases and their ability to sulfate luciferins, p-nitrophenol, and other small molecule substrates.**

Blank cells indicate missing or unknown data.

*Sulfates the luciferin coelenterazine at a different site compared to sea pansy sulfated luciferin, as shown in Figure S2.

**The ability to sulfate luciferin in VtST3 and VtST5 was supported by both the commercially available sulfotransferase assay and the luminescence assay.

*Supplemental References*

1. Copeland RA. 2000 *Enzymes: A Practical Introduction to Structure, Mechanism, and Data Analysis*. John Wiley & Sons.

2. Hempel N, Barnett A, Garnage N, Duggleby RG, Windmill KF, Martin JL, McManus ME. 2005 10 Human SULT1A SULTs. *Human cytosolic sulfotransferases* , 179.

3. Kakuta Y, Pedersen LG, Pedersen LC, Negishi M. 1998 Conserved structural motifs in the sulfotransferase family. *Trends Biochem. Sci.* **23**, 129–130.

4. Komatsu K, Driscoll WJ, Koh YC, Strott CA. 1994 A P-loop related motif (GxxGxxK) highly conserved in sulfotransferases is required for binding the activated sulfate donor. *Biochem. Biophys. Res. Commun.* **204**, 1178–1185.

5. Goodheart JA *et al.* 2020 Laboratory culture of the California Sea Firefly Vargula tsujii (Ostracoda: Cypridinidae): Developing a model system for the evolution of marine bioluminescence. *Sci. Rep.* **10**, 1–15.

6. Kushida A, Horie R, Hattori K, Hamamoto H, Sekimizu K, Tamura H. 2012 Xanthurenic acid is an endogenous substrate for the silkworm cytosolic sulfotransferase, bmST1. *J. Insect Physiol.* **58**, 83–88.

7. Bairam AF, Kermasha ZW, Liu M-C, Kurogi K, Yamamoto K. 2020 Functional analysis of novel sulfotransferases in the silkworm Bombyx mori. *Arch. Insect Biochem. Physiol.* **104**, e21671.

8. Yamamoto K, Yamada N, Endo S, Kurogi K, Sakakibara Y, Suiko M. 2022 Novel silkworm (Bombyx mori) sulfotransferase swSULT ST3 is involved in metabolism of polyphenols from mulberry leaves. *PLoS One* **17**, e0270804.

9. Hattori K, Inoue M, Inoue T, Arai H, Tamura H-O. 2006 A novel sulfotransferase abundantly expressed in the dauer larvae of Caenorhabditis elegans. *J. Biochem.* **139**, 355–362.

10. Tsoi C, Falany CN, Morgenstern R, Swedmark S. 2001 Molecular cloning, expression, and characterization of a canine sulfotransferase that is a human ST1B2 ortholog. *Arch. Biochem. Biophys.* **390**, 87–92.

11. Suiko M, Sakakibara Y, Liu M-Y, Yang Y-S, Liu M-C. 2005 Cytosolic Sulfotransferases and Environmental Estrogenic Chemicals. *J. Pestic. Sci.* **30**, 345–353.

12. Yasuda S, Kumar AP, Liu M-Y, Sakakibara Y, Suiko M, Chen L, Liu M-C. 2005 Identification of a novel thyroid hormone-sulfating cytosolic sulfotransferase, SULT1 ST5, from zebrafish. *FEBS J.* **272**, 3828–3837.

13. Yasuda S, Liu C-C, Takahashi S, Suiko M, Chen L, Snow R, Liu M-C. 2005

Identification of a novel estrogen-sulfating cytosolic SULT from zebrafish: molecular cloning, expression, characterization, and ontogeny study. *Biochem. Biophys. Res. Commun.* **330**, 219–225.

14. Liu T-A *et al.* 2008 Identification and characterization of two novel cytosolic sulfotransferases, SULT1 ST7 and SULT1 ST8, from zebrafish. *Aquat. Toxicol.* **89**, 94–102.

15. Mohammed YI *et al.* 2012 Identification and characterization of zebrafish SULT1 ST9, SULT3 ST4, and SULT3 ST5. *Aquat. Toxicol.* **112-113**, 11–18.

16. Yasuda S, Liu M-Y, Yang Y-S, Snow R, Takahashi S, Liu M-C. 2006 Identification of novel hydroxysteroid-sulfating cytosolic SULTs, SULT2 ST2 and SULT2 ST3, from zebrafish: Cloning, expression, characterization, and developmental expression. *Arch. Biochem. Biophys.* **455**, 1–9.

17. Yasuda T, Yasuda S, Williams FE, Liu M-Y, Sakakibara Y, Bhuiyan S, Snow R, Carter G, Liu M-C. 2008 Characterization and ontogenic study of novel steroid-sulfating SULT3 sulfotransferases from zebrafish. *Mol. Cell. Endocrinol.* **294**, 29–36.

18. Yasuda S *et al.* 2009 A novel hydroxysteroid-sulfating cytosolic sulfotransferase, SULT3 ST3, from zebrafish: identification, characterization, and ontogenic study. *Drug Metab. Lett.* **3**, 217–227.

19. Sugahara T, Liu CC, Govind Pai T, Liu MC. 2003 Molecular cloning, expression, and functional characterization of a novel zebrafish cytosolic sulfotransferase. *Biochem. Biophys. Res. Commun.* **300**, 725–730.

20. Hattori K, Motohashi N, Kobayashi I, Tohya T, Oikawa M, Tamura H-O. 2008 Cloning, expression, and characterization of cytosolic sulfotransferase isozymes from Drosophila melanogaster. *Biosci. Biotechnol. Biochem.* **72**, 540–547.

21. Wilson LA, Reyns GE, Darras VM, Coughtrie MWH. 2004 cDNA cloning, functional expression, and characterization of chicken sulfotransferases belonging to the SULT1B and SULT1C families. *Arch. Biochem. Biophys.* **428**, 64–72.

22. Cao H, Agarwal SK, Burnside J. 1999 Cloning and expression of a novel chicken sulfotransferase cDNA regulated by GH. *J. Endocrinol.* **160**, 491–500.

23. Sakakibara Y, Suiko M, Pai TG, Nakayama T, Takami Y, Katafuchi J, Liu MC. 2002 Highly conserved mouse and human brain sulfotransferases: molecular cloning, expression, and functional characterization. *Gene* **285**, 39–47.

24. Inouye S, Matsuda K, Nakamura M. 2023 Enzymatic sulfation of coelenterazine by human cytosolic aryl sulfotransferase SULT1A1: identification of coelenterazine C2-benzyl monosulfate by LC/ESI-TOF-MS. *Biochem. Biophys. Res. Commun.* **665**, 133–140.

25. Gamage N, Barnett A, Hempel N, Duggleby RG, Windmill KF, Martin JL, McManus ME. 2005 Human Sulfotransferases and Their Role in Chemical Metabolism. *Toxicol. Sci.* **90**, 5–22.

26. Wang J, Falany JL, Falany CN. 1998 Expression and characterization of a novel thyroid hormone-sulfating form of cytosolic sulfotransferase from human liver. *Mol. Pharmacol.* **53**, 274–282.

27. Kurogi K, Shimohira T, Kouriki-Nagatomo H, Zhang G, Miller ER, Sakakibara Y, Suiko M, Liu M-C. 2017 Human Cytosolic Sulphotransferase SULT1C3: genomic analysis and functional characterization of splice variant SULT1C3a and SULT1C3d. *J. Biochem.* **162**, 403–414.

28. Hempel N, Barnett AC, Bolton-Grob RM, Liyou NE, McManus ME. 2000 Site-directed mutagenesis of the substrate-binding cleft of human estrogen sulfotransferase. *Biochem. Biophys. Res. Commun.* **276**, 224–230.

29. Her C, Wood TC, Eichler EE, Mohrenweiser HW, Ramagli LS, Siciliano MJ, Weinshilboum RM. 1998 Human hydroxysteroid sulfotransferase SULT2B1: two enzymes encoded by a single chromosome 19 gene. *Genomics* **53**, 284–295.

30. Sun Y, Machalz D, Wolber G, Parr MK, Bureik M. 2020 Functional Expression of All Human Sulfotransferases in Fission Yeast, Assay Development, and Structural Models for Isoforms SULT4A1 and SULT6B1. *Biomolecules* **10**. (doi:10.3390/biom10111517)

31. Pichu S, Yalcin EB, Ribeiro JM, King RS, Mather TN. 2011 Molecular characterization of novel sulfotransferases from the tick, Ixodes scapularis. *BMC Biochem.* **12**, 32.

32. Takahashi S, Sakakibara Y, Mishiro E, Kouriki H, Nobe R, Kurogi K, Yasuda S, Liu M-C, Suiko M. 2009 Molecular Cloning, Expression and Characterization of A Novel Mouse SULT6 Cytosolic Sulfotransferase. *J. Biochem.* **146**, 399–405.

33. Sakakibara Y, Yanagisawa K, Takami Y, Nakayama T, Suiko M, Liu MC. 1998 Molecular cloning, expression, and functional characterization of novel mouse sulfotransferases. *Biochem. Biophys. Res. Commun.* **247**, 681–686.

34. Saeki Y, Sakakibara Y, Araki Y, Yanagisawa K, Suiko M, Nakajima H, Liu MC. 1998 Molecular cloning, expression, and characterization of a novel mouse liver SULT1B1 sulfotransferase. *J. Biochem.* **124**, 55–64.

35. Tzertzinis G, Baker B, Benner J, Brown E, Corrêa IR Jr, Ettwiller L, McClung C, Schildkraut I. 2022 Coelenterazine sulfotransferase from Renilla muelleri. *PLoS One* **17**, e0276315.

36. Song WC, Moore R, McLachlan JA, Negishi M. 1995 Molecular characterization of a testis-specific estrogen sulfotransferase and aberrant liver expression in obese

and diabetogenic C57BL/KsJ-db/db mice. *Endocrinology* **136**, 2477–2484.

37. Shimada M, Terazawa R, Kamiyama Y, Honma W, Nagata K, Yamazoe Y. 2004 Unique properties of a renal sulfotransferase, St1d1, in dopamine metabolism. *J. Pharmacol. Exp. Ther.* **310**, 808–814.

38. Fallon TR, Li F-S, Vicent MA, Weng J-K. 2016 Sulfoluciferin is Biosynthesized by a Specialized Luciferin Sulfotransferase in Fireflies. *Biochemistry* **55**, 3341–3344.
